# Calcareous sponge cell atlas provides support to homology between sponge and eumetazoan body plans

**DOI:** 10.64898/2026.02.26.708390

**Authors:** Di Pan, Dinithi Rajapaksha, Cüneyt Caglar, Rachel Rathjen, Marcin Adamski, Maja Adamska

## Abstract

Sponges are widely recognized as important model organisms for studying animal evolution, due to their phylogenetic position at the base of the animal tree of life, as well as similarities to the nearest animal relatives, the choanoflagellates. A critical aspect of animal evolution concerns the origin of germ layers, the embryonic structures which give rise to all tissues and organs of animal bodies. Haeckel’s hypothesis suggested a germ layer homology between sponges and corals, and thus all eumetazoans (complex animals including cnidarians and bilaterians). According to this hypothesis, sponge choanoderm (composed of the feeding cells, choanocytes) and sponge pinacoderm (the outer epithelium) would be homologous to eumetazoan endoderm (from which the digestive system originates) and the ectoderm (giving rise to the epidermis), respectively. We addressed this hypothesis comparing tissue-specific transcriptomes derived from single-cell transcriptome datasets of sponges and cnidarians. We have sequenced single cell transcriptomes of Australian calcareous sponge, *Sycon capricorn*, and identified its cell types using a combination of in silico annotation of the cell clusters and in situ hybridization with marker genes. Single-cell transcriptome datasets for two demosponge species and two cnidarian species were extracted from recent literature. Homology was assessed using the SAMap algorithm, which has been designed to identify homologous cell types across vast evolutionary distances by detection of shared expression profiles. Our results are fully consistent with Haeckel’s hypothesis, supporting homology between the innermost layers of sponges and cnidarians (choanoderm and endoderm/gastrodermis) as well as the outermost layers of sponges and cnidarians (pinacoderm and ectoderm/epidermis). Thus, sponge body plan appears to represent an intermediate step between single cell protists (choanoflagellates) and complex animals, rather than being independent experiment in animal multicellularity as suggested by alternative hypotheses.

## Introduction

Multicellular animals (Metazoa) most likely evolved from colonial protists resembling modern choanoflagellates, capable of temporal cell differentiation involving epithelial-like and amoeboid cells (King 2004; Maldonado 2004; Mikhailov et al. 2009; Brunet et al. 2021). The subsequent grades of organization of the ancestors along the way leading generation of the diversity of extant animals (Figure 1a) remain an area of active investigation and heated debates (e.g. King & Rokas 2017).

**Figure 1.**
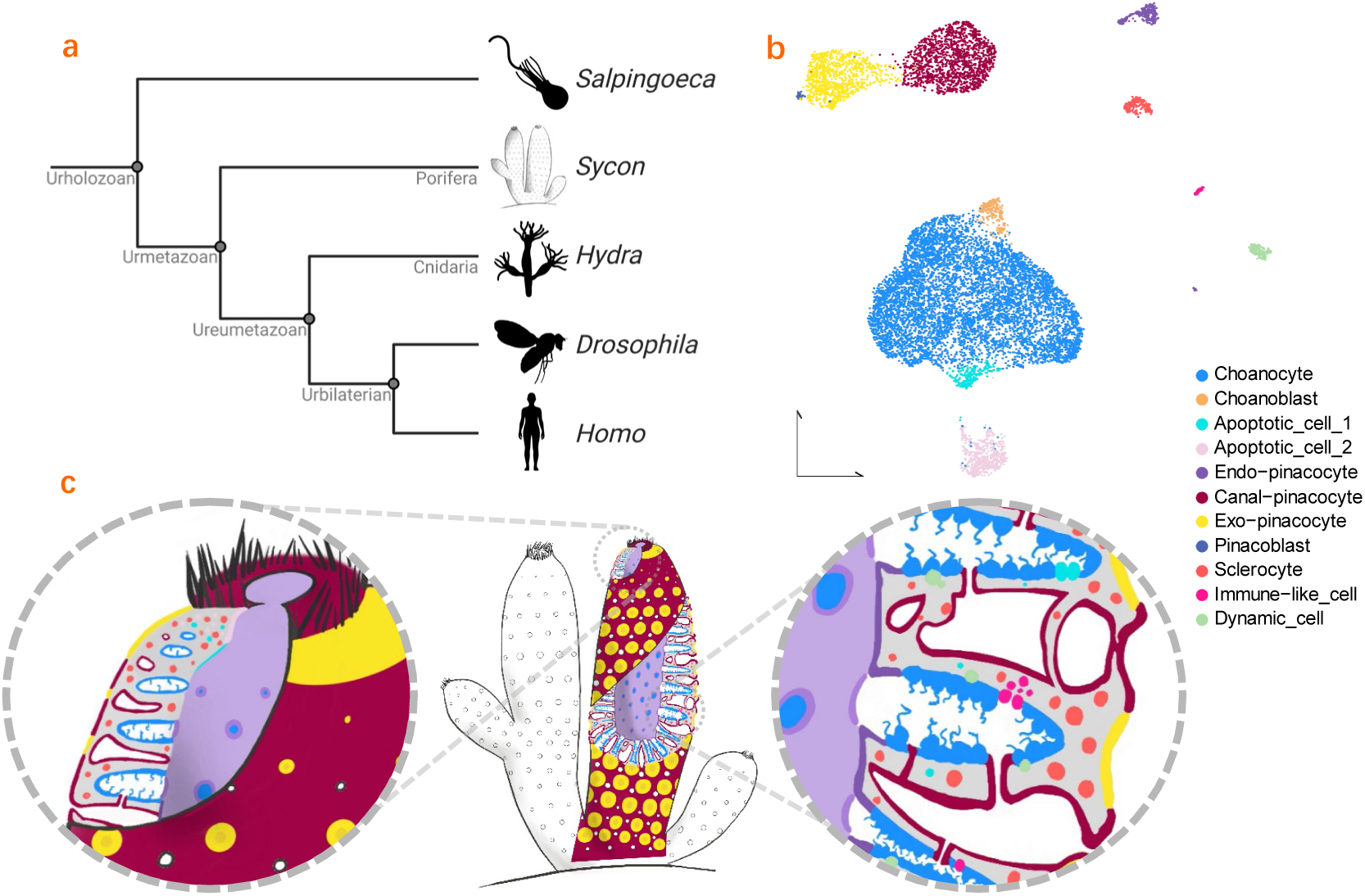
*Sycon capricorn* and its cell types defined by single-cell RNA sequencing presented in this study. **a**, Phylogenetic position of *S. capricorn* in relation to hypothetical ancestors of the major animal clades. **b**, UMAP plot of 10,747 *S. capricorn* cells coloured by 11 clusters. Each colour represents a cell type; **c**, Morphology of *S. capricorn*, showing localisation of the cell types identified by scRNA analysis.

Majority of multicellular animals belong to the clade Bilateria, which are a monophyletic group characterized by bilateral symmetry, digestive system usually in the form of a through gut, as well as diverse cell types including neurons and muscles. In Bilaterians, tissues and organs are derived from three embryonic germ layers: ectoderm (giving rise to epidermis and the nervous system), mesoderm (producing, among others, muscle), and endoderm (producing the digestive system and associated organs). These germ layers are formed during gastrulation, a process that occurs early in embryonic development and involves the movement and rearrangement of cells (Solnica-Krezel & Sepich 2012).

The sister group to Bilaterians, the Cnidarians, also undergo gastrulation during their embryonic development, but their bodies are generally composed of only two layers, epidermis and gastrodermis, in fact often called, especially in reference to the freshwater *Hydra* species, ectoderm and endoderm. Traditionally, mesoderm has been considered to be bilaterian evolutionary novelty, derived from endoderm present in the last common ancestor of cnidarians and bilaterians (the Ureumetazoan), although recent studies suggest more complex relationships between cnidarian and bilaterian tissue layers (Technau & Scholz 2003; Martindale, Pang & Finnerty 2004; Genikhovich & Technau 2017; Steinmetz et al. 2017; Haillot et al. 2025). Nonetheless, existence of Eumetazoan ancestor possessing nerves and muscles, and body consisting of the inner, digestive, endodermally derived layer, and an outer, ectodermally derived layer serving protective role does not appear to be contested.

On the other hand, the nature of the Urmetazoan – the ancestor of all multicellular animals, including sponges, ctenophores and placozoans in addition to cnidarians and bilaterians, remains elusive, confounded by conflicting phylogenies, alternatively placing sponges or ctenophores as the sister group to all remaining metazoans (Schultz et al. 2023; Copley 2025). To make matters more complicated, the ctenophore complex body plans and diverse cell types might have evolved independently from those in the ancestor of cnidarians and bilaterians (Ryan et al. 2013; Moroz et al. 2014), while the placozoans might be secondarily simplified. With these caveats in mind, here we focus on attempting to gain insight into the body plan and cell types present in the last common ancestor of modern sponges and eumetazoans, remaining agnostic to the phylogenetic position (and thus, evolutionary history of body plans) of ctenophores and placozoans.

We are particularly interested in understanding whether germ layers were already present in the last common ancestor of sponges and eumetazoans. While not widely accepted (Leininger et al. 2014 vs Nakanishi, Sogabe & Degnan 2014), one of the earliest theories about the origin of germ layers posited that germ layers evolved from the ancestral body plan of sponges (Haeckel 1874). Haeckel (1870) viewed the similarities in structure and function between sponges (particularly calcareous sponges) and cnidarian polyps as evidence for a direct evolutionary link between the inner tissue layer of eumetazoans (endoderm) and the inner layer of sponges (choanoderm), as well as between the outer tissue layer of eumetazoans (ectoderm) and the outer layer of sponges (pinacoderm).

Here, we use calcareous sponge *Sycon capricorn* (Woerheide & Hooper 2003) to generate single cell atlas (Figure 1b-c) and, by comparison with published singe cell datasets, detect homologous cell types between sponges and cnidarians.

## Results and Discussion

### *S. capricorn* is composed of 11 distinct cell types

Calcareous sponges exhibit exceptional regenerative capacity, including ability to undergo complete regeneration from dissociated cells, as observed in syconoid calcareous sponges like *Sycon raphanus*, *Sycon lingua*, *Sycon ciliatum* (Huxley 1912; Soubigou et al. 2020; Caglar et al. 2021; Ereskovsky et al. 2021). *S. capricorn* cells can rapidly re-aggregate within minutes post-dissociation and produce juvenile-like entities within weeks (figure S1). In this study, we employed the ACME (acetic acid with methanol, García-Castro et al. 2021) fixation method combined with fluorescence-activated cell sorting (FACS) (figure S2) to produce single cell atlas for *S. capricorn*. Following the consolidation and filtration of 10,747 cells from four pooled adult sample replicates (figure S3), 11 distinct cell types were identified and characterized (Figure 1b, and figure S4-S8).

To annotate the identified cell types, we used a combination of in situ hybridization (ISH), GO-enrichment and analysis of cell trajectories. ISH, with RNA probes based on marker genes identified for each cell cluster, allowed mapping of cell clusters to cells in sponge tissues. This approach allowed us to annotate the major identified clusters as known sponge cell types with characteristic morphologies and locations (choanocytes, pinacocytes and sclerocytes). Additional analyses, such as GO-term enrichment of cluster markers, allowed annotation of cell clusters in cases where morphology or position of cells were not sufficient to identify a cell type.

Choanocytes, which have the function of filter-feeding and stemness in calcareous sponges, have been detected in a high proportion, as expected (Leys & Eerkes-Medrano 2006; Adamska 2018; Funayama 2018). Choanocytes form chambers in choanoderm and have a single flagellum surrounded by microvillous collars (Figure 2, and figure S10). One cell type similar in nature to the choanocyte cluster is the choanoblast, distinguished by its exclusive expression of cell cycle-related genes (figure S9), such as *PCNA* (Schoenenberger et al. 2015), *CyclinB* (Takizawa & Morgan 2000), and *Ki67* (Uxa et al. 2021). Choanoblasts are dispersed within the choanocyte chambers and exhibit morphological resemblance to choanocytes (Figure 2, and figure S10). Another cell type close to the choanocyte cluster was called apoptotic cell 1 (see below for explanation of the name). They primarily localize within the basal region of the osculum’s inner wall as well as in sporadic cell clusters within the choanocyte chambers and exist in small amounts in mesohyl (Figure 2, and figure S11). The outcomes of the gene ontology (GO) analysis indicated their involvement in early apoptotic processes (Figure 3). Apoptotic cell 1 specifically expresses *p53*, which can induce cell cycle arrest and apoptosis (figure S11c) (Ozaki & Nakagawara 2011). In the UMAP analysis (Figure 1b), a cluster that was distinct from the aforementioned three cell type clusters and exhibited a ‘tailing’ pattern was designated as apoptotic cell 2. They localize adjacent to apoptotic cell 1, primarily in the up region of the inner wall in the osculum rather than below (Figure 2). They can also be observed in the mesohyl and choanocyte chambers (figure S11). The outcomes of the GO analysis indicated their involvement in late apoptotic processes (Figure 3). As predicted, apoptotic cell 2 specifically expressed genes associated with apoptosis and DNA repair (figure S11c), such as *TRAF3* (Zhou et al. 2021), *PNKP* (Weinfeld et al. 2011), and *TNKS1* (Kim 2018).

**Figure 2.**
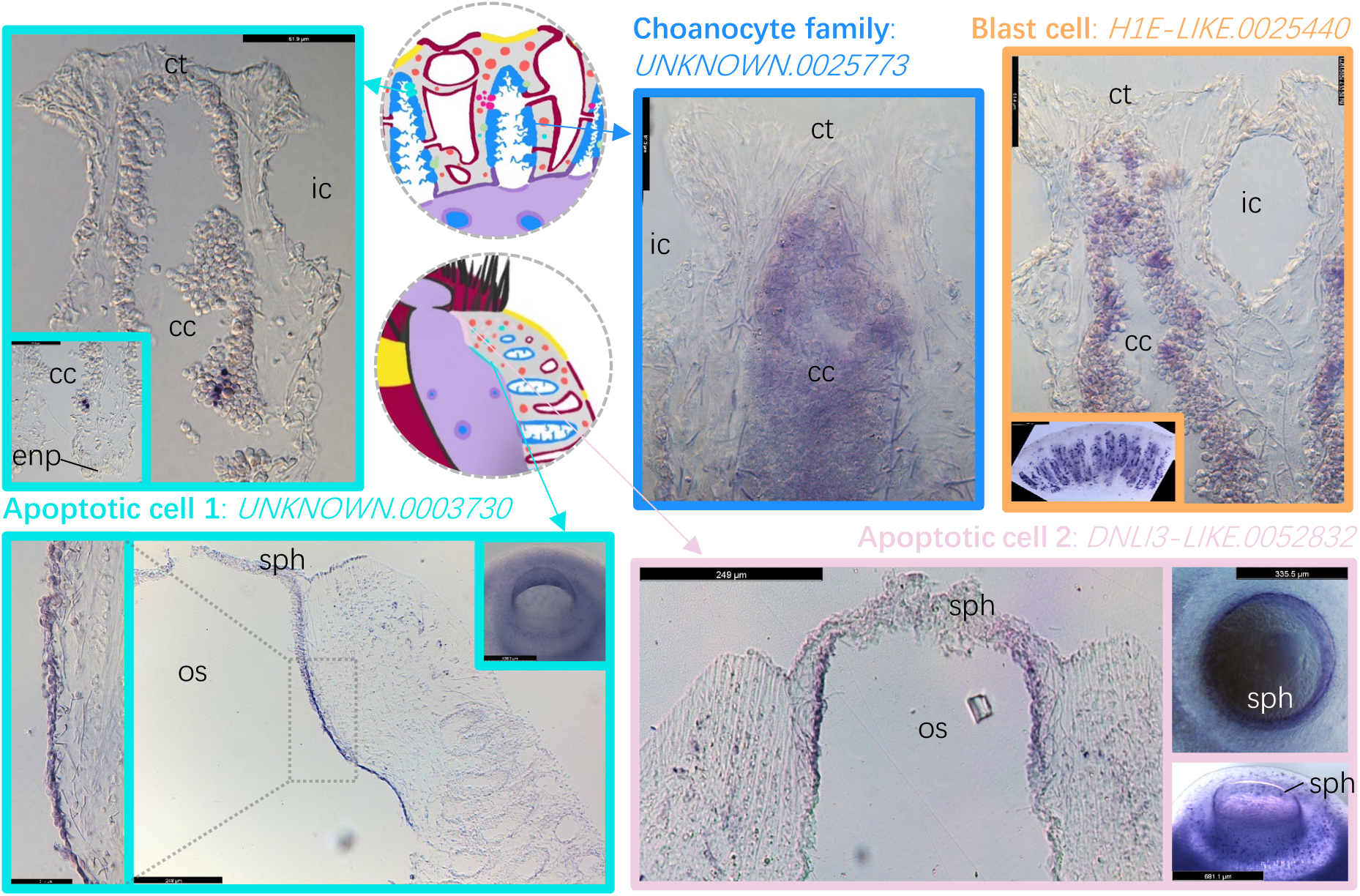
Localisation of cell types in choanocyte family. The names of the cell types and the marker gene used are marked in the figure. The overall positioning of the **choanocyte family** is marked with a blue border. The choanocyte family includes four cell types, including choanocytes, mainly concentrated in the choanocyte chamber; The **blast cells**, which include choanoblast and pinacoblast, are labelled with the same marker and are outlined with an orange border. The images show that in addition to the choanoblasts that appear in the chambers, cells in other locations are also labelled, such as among the exo-pinacocytes, which are likely the pinacoblast; The remaining two border colours, cyan and pink, represent the **apoptotic cell 1** and **2**. They are distributed in the upper and lower parts of the inner wall of the osculum and in small quantities in other areas, such as the osculum mesohyl. cc - choanocyte chamber; ct - chamber tip; enp - endo-pinacocyte; ic - incurrent canal; os - osculum; sph - sphincter.

**Figure 3.**
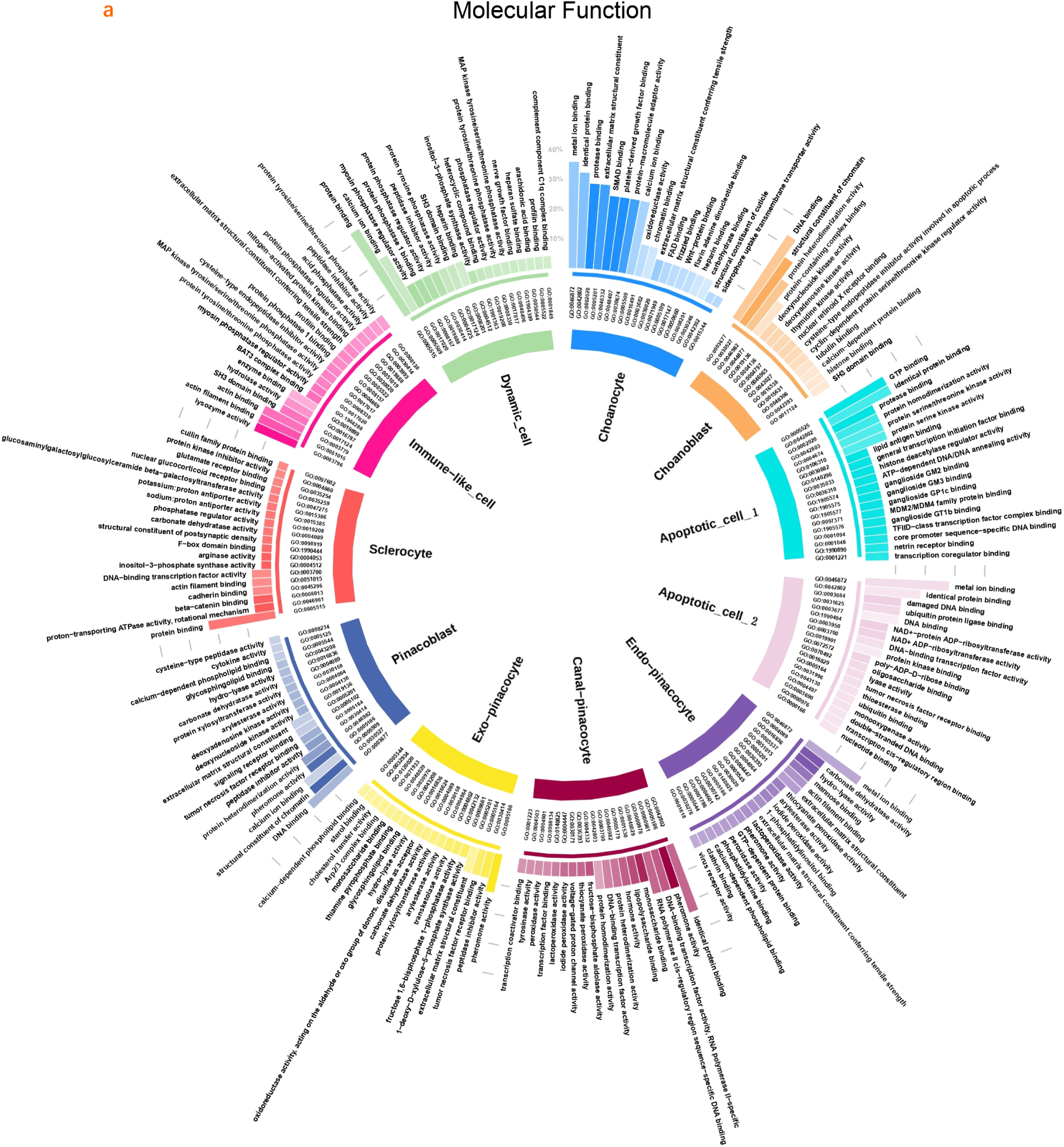
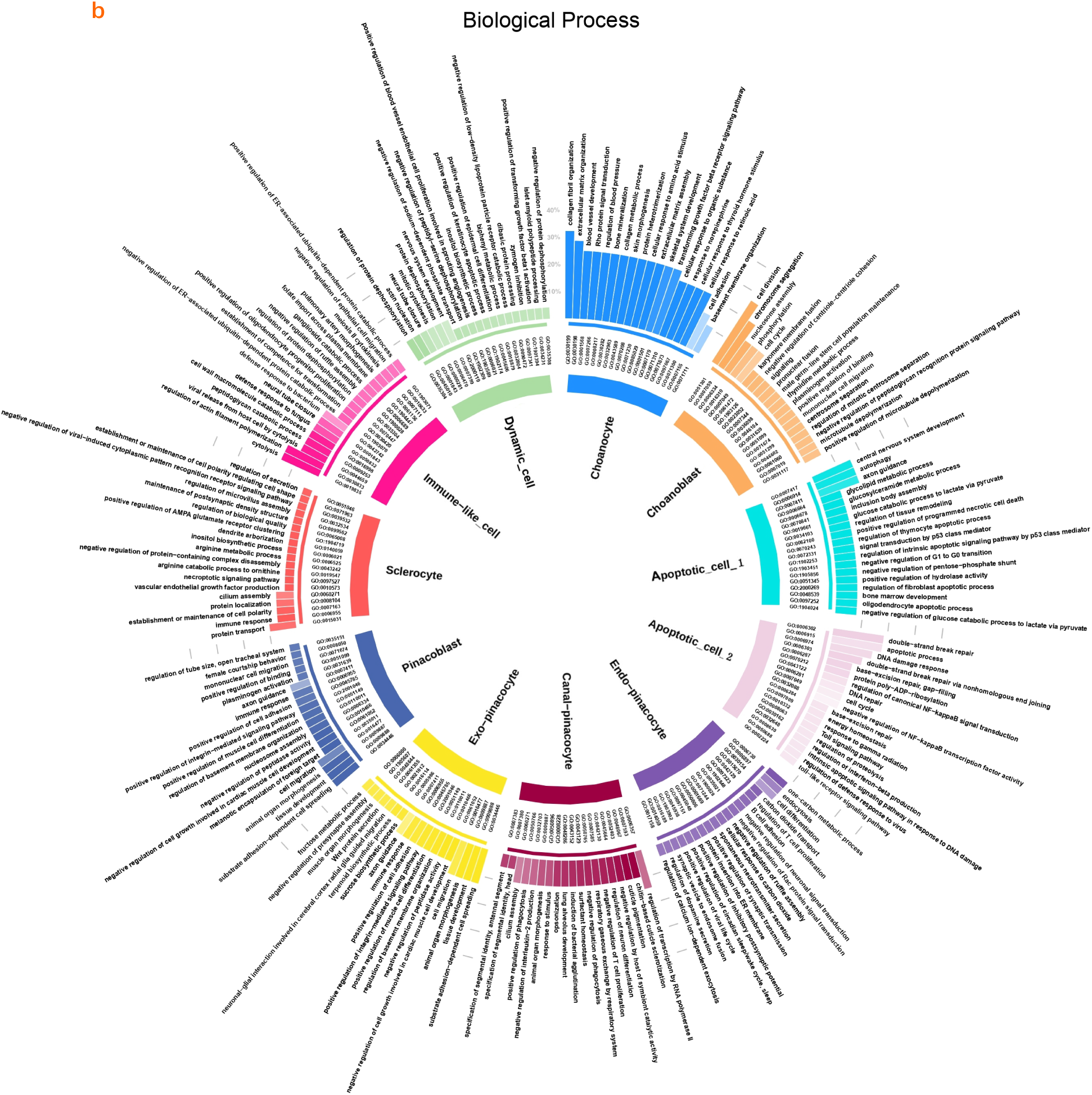
GO-term enrichment of each cell type. Molecular function (**a**) and biological process (**b**) GO-term enrichment. Blast cells were highly enriched for cell proliferation-related events. Apoptotic cells 1 and 2 were highly enriched for early and late events related to DNA damage and repair. Immune-like cells were highly enriched for immunity-related events. The height of each bar represents the gene ratio. The darker colour represents the lower p-value. More details are presented in figure S7-S8.

In addition to choanocytes, pinacocytes, which usually have a flat shape, are another abundant epithelial cell type in sponges (Adamska 2018). On the body surface of the *S. capricorn*, exo-pinacocytes appear at the top of the choanocyte chamber, which we refer to as the chamber tip (Figure 4). From a macroscopic perspective, the distribution area of exo-pinacocytes appears as regularly arranged patches. There is also a large number of exo-pinacocytes distributed on the surface of the osculum (figure S12). Canal-pinacocytes are present on the body surface of the sponge in complementary forms. Between the chambers are channels for the circulation of seawater, which are composed of canal-pinacocytes. From the surface, the canal-pinacocytes are distributed in a grid pattern, tightly framing the chamber tips where the exo-pinacocytes are located (Figure 4, and figure S13). The sphincter that controls the rate at which seawater flows out of the osculum is also composed of canal-pinacocytes. The cells in the cluster adjacent to exo-pinacocytes on the opposite side of canal-pinacocytes on UMAP are called pinacoblasts. Pinacoblasts and choanoblasts highly express many genes related to the cell cycle (Figure 2, and figure S10). They are also enriched with many proliferation-related GO terms (Figure 3), so they are collectively called ‘blast cells’ here. They are challenging to locate individually because they are rare. The epithelial cells of the spongocoel between the chamber openings are endo-pinacocytes (Figure 4, figure S13). The endo-pinacocytes extend upward along the spongocoel and border apoptotic cell 1 near the osculum.

**Figure 4.**
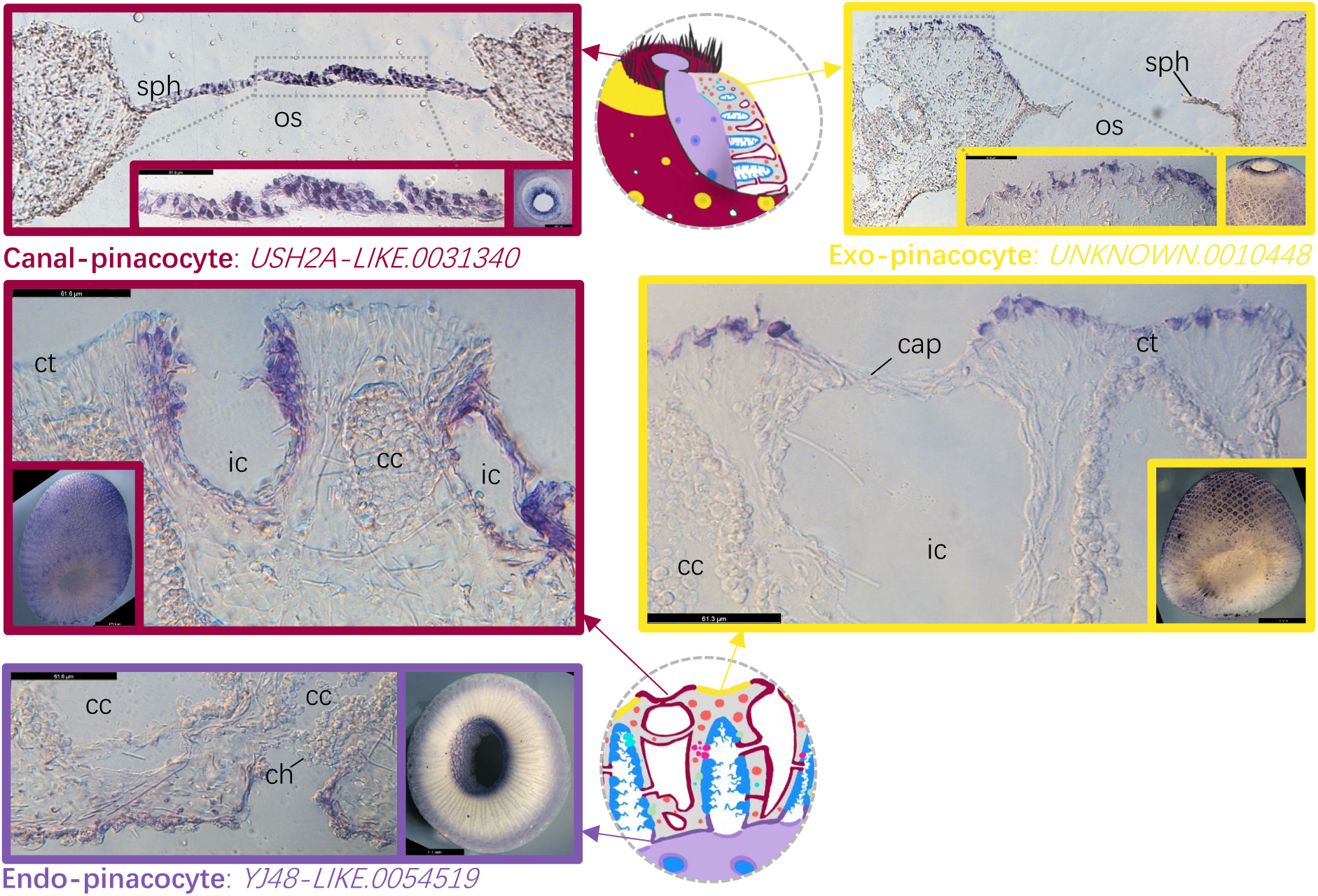
Localisation of cell types in pinacocyte family. Each colour of the image border represents a cell type. The name of the cell type and the marker gene used are marked. Dark red represents **canal-pinacocytes** found on the channel surface and the oscular sphincter. Yellow represents **exo-pinacocytes**, which were found on the chamber tip surface and osculum surface. Purple represents **endo-pinacocytes**, which were found on the spongocoel surface. cap - canal-pinacocyte; cc - choanocyte chamber; ch - choanocyte; ct chamber tip; ic - incurrent canal; os - osculum; sph - sphincter.

Sclerocytes are amoeboid cells and are responsible for producing spicules (i.e., skeletal elements) in the mesohyl layer of calcareous sponges (Voigt et al. 2014; Voigt et al. 2017). Sclerocytes usually exist in clusters and can be found anywhere in the mesohyl, with particular concentrations at the chamber tip (Figure 5, and figure S14). Therefore, from the perspective of the epidermis, its distribution is similar to that of the reticular canal-pinacocytes but is located below the surface. Immune cells, also known as grey cells, are another type of cell found in sponge mesohyl (Yin & Humphreys 1996; Bergquist 1998). Upon foreign pathogen invasion, grey cells congregate near the infected cells, immobilizing and preventing further spread. Should the invasion persist, these cells release toxins, eradicating all cells in the vicinity to ensure defense. A type of cell in *S. capricorn* was highly enriched with immune-related terms in GO analysis (Figure 3) and was therefore called immune-like cell. These cells were often found in the form of clusters near the chamber surface (Figure 5, and figure S15). The last cell cluster to be annotated is referred to as ‘dynamic cell’ due to the apparently highest dynamics of gene expression revealed through RNA velocity analysis (Figure 6a). They were usually found in the mesohyl area next to the choanocytes (Figure 5, and figure S15).

**Figure 5.**
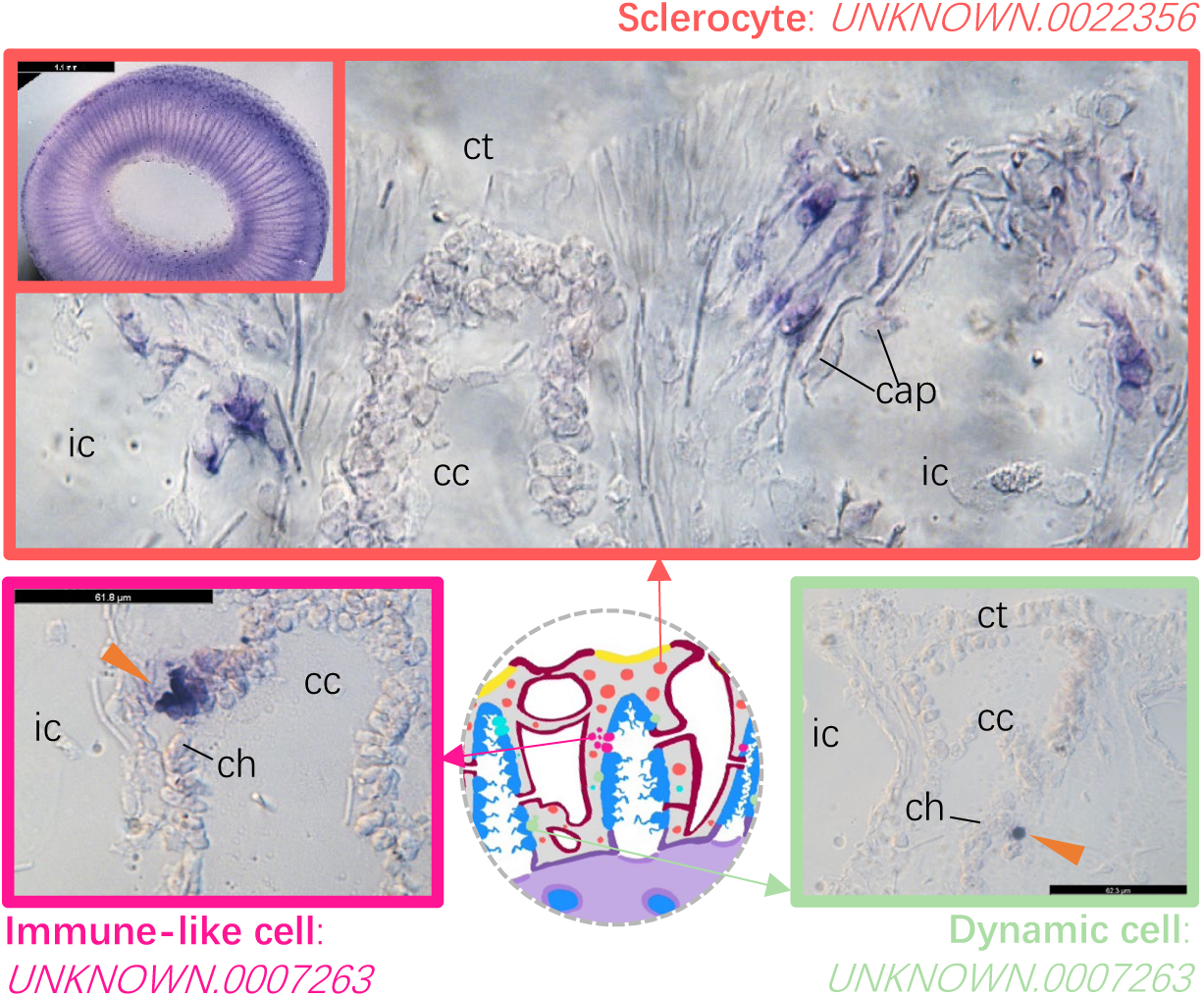
Localisation of mesenchymal cells. Each colour of the image border represents a cell type. The name of the cell type and the marker gene used are marked in the figure. Orange represents **sclerocytes**, which are concentrated in the mesohyl at the chamber tip and often exist in clusters. Pink represents **immune-like cells**, which can be found in the mesohyl close to the chamber surface and exist in clusters (as indicated by the arrow). Green represents **dynamic cells**, which can be found in the mesohyl adjacent to the choanocyte (as indicated by the arrow). cap - canal-pinacocyte; cc - choanocyte chamber; ch - choanocyte; ct - chamber tip; ic - incurrent canal.

**Figure 6.**
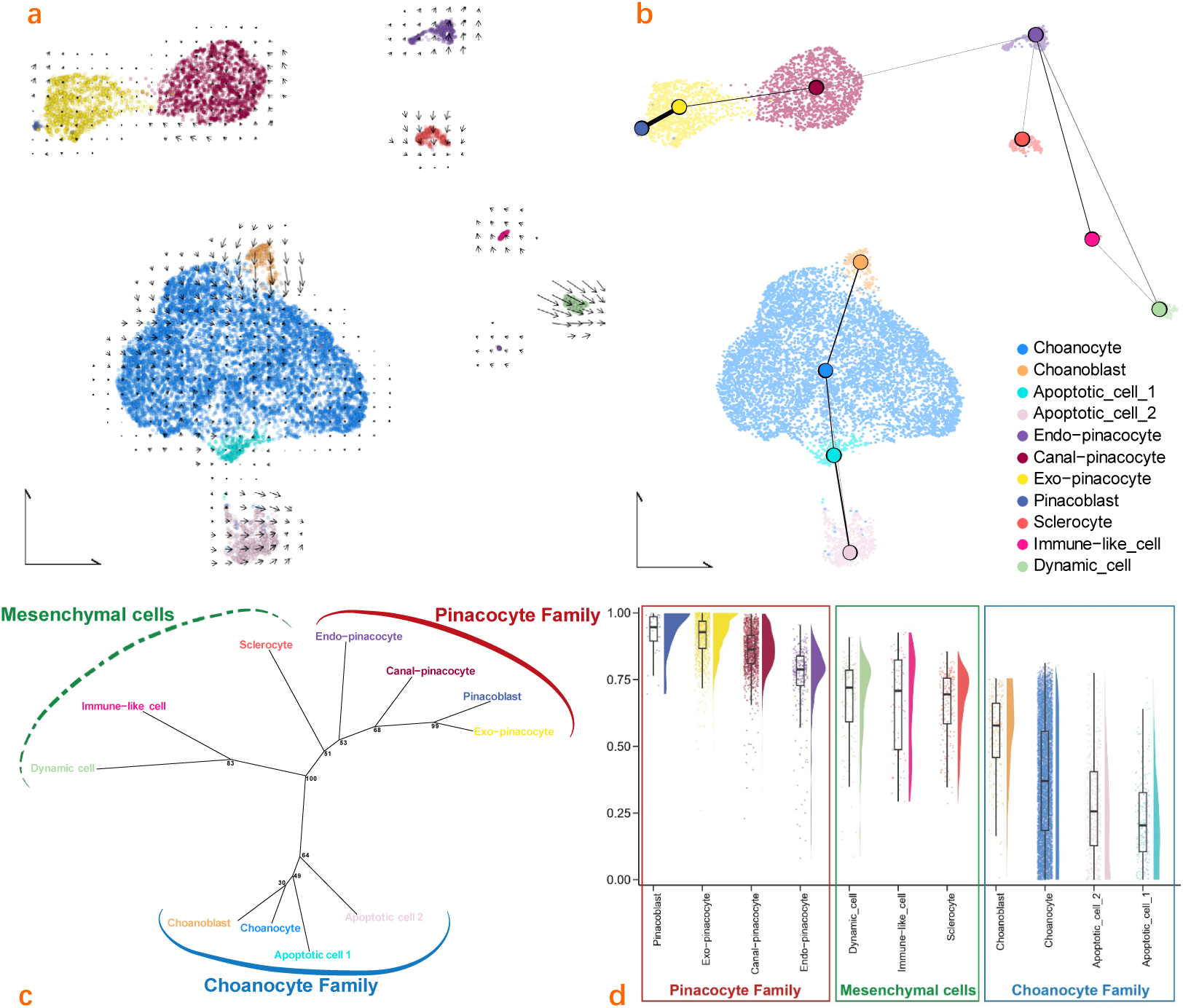
*S. capricorn* cell type trajectory and families. **a**, RNA velocity plot showing differences in RNA splicing rates between cells; **b**, PAGA connectivity graph exhibiting connection strength (colour depth and edge thickness); **c**, Cell type phylogeny with bootstrap support; **d**, CytoTRACE plot showing decreasing transcriptional diversity among cell types from left to right in each family.

### Interrelationships of cell types in *S. capricorn*

RNA velocity analysis was conducted to forecast the future state of each individual cell (Figure 6a). In the UMAP visualization, the choanoblast cluster appears as an initial point dynamically transitioning towards the adjacent choanocyte cluster. It appears that choanocytes progress towards becoming apoptotic cell 1, and while no significant direct transition from apoptotic cell 1 to apoptotic cell 2 is evident, apoptotic cells 2 maintain distinctive identity without transitioning into other cell types. In contrast to choanoblasts, pinacoblasts exhibit a substantially lower initiation rate as starting points. Moreover, alongside the previously discussed dynamic cell populations, the conversion rates of pinacocytes and mesohyl cells are notably minimal, i.e., cell identities are stable.

We further explored developmental relationships using the partition-based graph abstraction (PAGA) connectivity graph (Figure 6b), which assesses the strength of connections between cell types. This analysis revealed strong connections between choanoblasts, choanocytes and the two apoptotic cells clusters, as well as pinacoblasts, exo-pinacocytes and canal-pinacocytes. Interestingly, the endo-pinacocyte cluster appears connected to both pinacocytes and mesohyl cells. We have produced a neighbour-joining tree representing relationships between *S. capricorn* cells, using methods modified from Musser et al. (2021) based on bootstrapping and the high variant genes (HVGs) of each cell type (Figure 6c). The findings were in alignment with the PAGA outcomes, whereby the four clusters associated with choanocytes coalesced into a distinct cluster denoted as the choanocyte family. Similarly, the clusters linked to pinacocytes aggregated into a cluster identified as the pinacocyte family, residing in close proximity to mesenchymal cells, with endo-pinacocyte serving as the connecting point. We further compared the developmental potential of each cell type and found that they were still arranged in the above three cell families (Figure 6d). Blast cells were always the cell type with the greatest differentiation potential in the choanocyte family and pinacocyte family.

Our results are consistent with the widely accepted notion of self-renewal potential of choanoblasts, which are likely to be multipotent stem cells in calcareous sponges. *PCNA*, *CyclinB*, and *Ki67* are associated with different phases of the cell cycle, from early to late (Kumar et al. 2009; Kousholt, Menzel & Sørensen 2012; Uxa et al. 2021). Different cells of choanoblasts highly express the above genes (figure S10), which further suggests the self-renewal potential of choanoblasts. Our trajectory analysis showed a clear cell fate that choanoblasts likely proliferate and differentiate into choanocytes and then turn into apoptotic cells. It is noteworthy that the conversion rate between apoptotic cells 1 and 2 is slow; and once cells are converted to apoptotic cell 2 successfully, their RNA velocity accelerates, which is consistent with the different stages of apoptosis. Notably, the slow conversion rate between apoptotic cells 1 and 2 and the accelerated RNA velocity once converted to apoptotic cell 2 is consistent with a cumulative initial phase of initiation or stimulation of apoptotic cell death and a later phase of active programmed cell death that occurs rapidly when events become irreversible (Best et al. 1999; Elmore 2007). Strikingly, the apoptotic cells of *S. capricorn* are concentrated in the osculum, and the cells that progress further into apoptosis are closer to the exhalant opening. We have found (Figure 2) that choanoblasts are distributed uniformly across the choanocyte chambers. Thus, we hypothesize that renewal of the choanoderm relies on production of new cells through the chambers with constant flow of cells pushed towards the osculum by the increased surface area, with choanocytes entering the oscular chimney activating the apoptotic pathway and likely being shed once they reach the opening. This process would be akin to renewal of the intestinal epithelium in vertebrates, and both epithelial layers in *Hydra*, where cells are produced by proliferation mid-body and shed at the terminal ends (Campbell 1967).

The links between identified cell types are further highlighted by visualisation of expression of selected developmental regulatory genes, including signalling molecules and transcription factors, which are strikingly co-expressed across cell families (figure S16). For example, pinacocyte family cells share expression of multiple *Wnt* and *Tgf-beta* genes (figure S16a), while the choanocyte family (including choanoblasts, choanocytes and both types of apoptotic cells) express a range of transcription factors associated with endoderm and mesoderm development across eumetazoans, including *Gata* and *Brachyury* (figure S16b) (Technau & Scholz 2003; Martindale, Pang & Finnerty 2004).

### Cell type homologies between two divergent sponge lineages

Calcareous sponges and demosponges have been evolving independently for approximately 600 million years and have been shown to have retained different subsets of ancestral developmental regulatory genes, including transcription factors (Fortunato, Adamski and Adamska 2015). The two lineages differ morphologically, with demosponges displaying leuconoid body plan organization, where choanocyte chambers are embedded in a thick, archaeocyte (stem cell) rich mesohyl layer, while calcareous sponges are can be asconoid (two epithelial layers of pinacoderm and choanoderm with narrow mesohyl layer); syconoid (addition of endo-pinacocytes lining the central atrium, various thickness of mesohyl) or leuconoid. In contrast to demosponges, not all calcareous sponges have archeocytes (reviewed by Adamska 2018), and consistently with this notion, our analysis has not revealed archeocytes in *S. capricorn*.

We were interested in revealing homologies between sponge cell types, both to investigate relationships between cell types evolving separately across large time scales, but also to confirm that our dataset and the applied methodology are capable of identifying relationships which are assumed to be true (for example, calcareous and demosponge pinacocytes) or are considered likely in light of the biology of the species (calcareous sponge choanocytes and demosponge archaeocytes). We have used previously published single cell datasets developed for demosponges: *Amphimedon queenslandica* (Sebé-Pedrós et al. 2018) and *Spongilla lacustris* (Musser et al. 2021), to compare *S. capricorn* cells using SAMap (Figure 7a). With relatively stringent criteria for detecting homologous cell relationships, not all cell types identified in the *S. capricorn* single cell atlas have been found to map to the demosponge cells. However, where relationships are found, they are in line with our knowledge of sponge biology, highlighting robustness of the methodology used.

**Figure 7.**
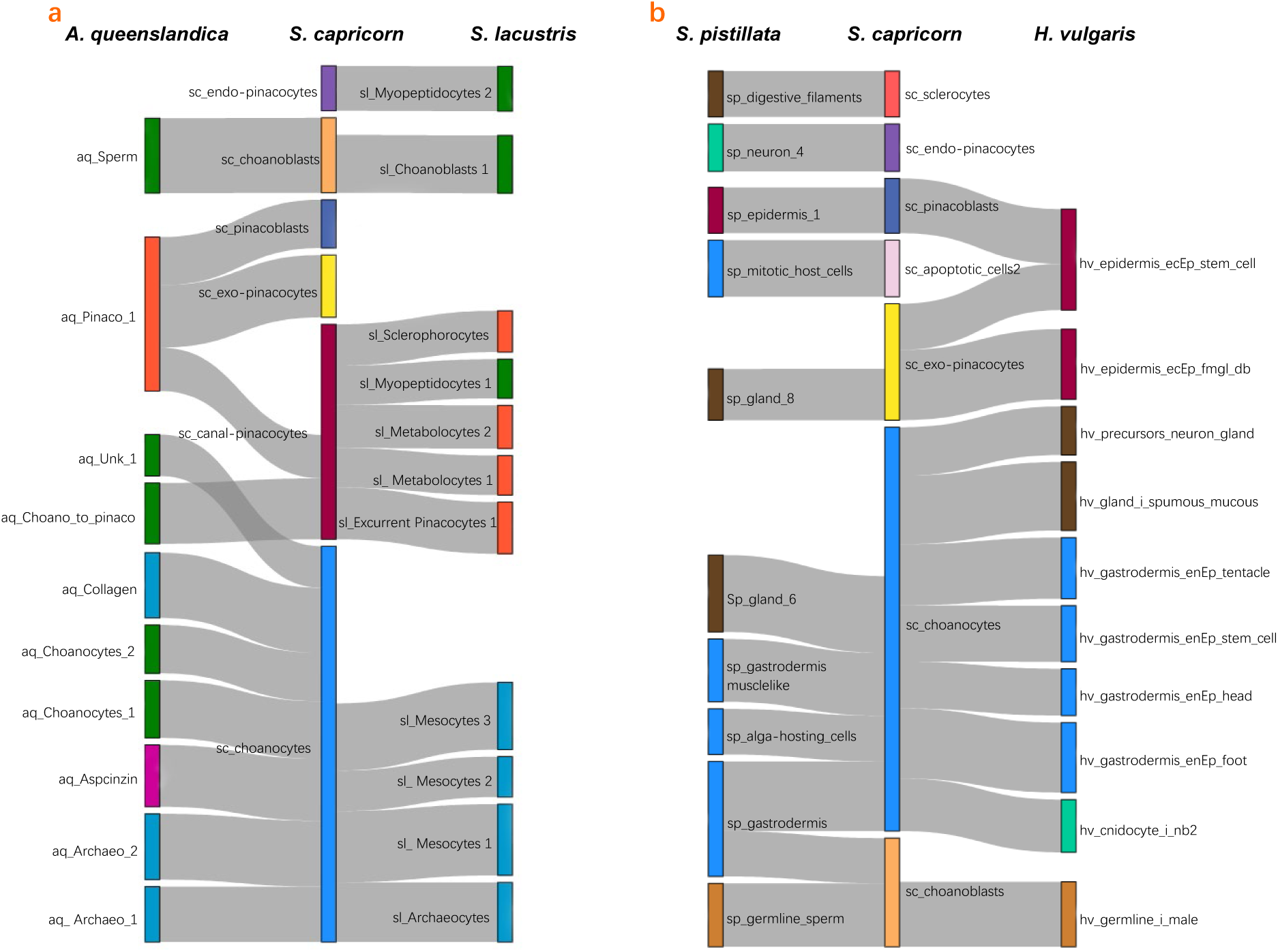
Sankey plots of single-cell SAMap alignment of *S. capricorn* with (a) sponges, *Amphimedon queenslandica* and *Spongilla lacustris*, and (b) cnidarians, *Stylophora pistillata* (coral) and *Hydra vulgaris* (freshwater hydrozoan). Each column in the plot represents cells from one species (name marked at the bottom. Each coloured rectangle represents a cell type. Except for *S. capricorn*, which uses the cell types and their colours as used throughout this study, the cell types of other species are divided into five categories and marked with different colours: Gastrodermis and related cell types: blue; Epidermis and related cell types: red; Neuronal cell types: green; Germline: orange; Other cell types: brown. The length of the rectangle represents the number of cells in it. The grey strips between columns represent the connection between cell types.

As expected, the exo-pinacocyte family cells from *Sycon* (pinacoblasts, exo-pinacocytes and canal-pinacocytes) mapped to pinacocytes and cells annotated as ‘choano-to-pinacocytes’ in *Amphimedon*, and to broad pinacocyte family (including pinacocytes, sclerophorocytes, and metabolocytes) in *Spongilla*. Interestingly, *Sycon* endo-pinacocytes and a fraction of canal-pinacocytes mapped to *Spongilla* myopeptidocytes, a cell type recently described in *Spongilla* (medium-sized mesenchymal cells with prominent vacuoles and long projections forming cellular network) (Musser et al. 2021). In line with our observation that *Sycon* endo-pinacocytes show links to both pinacocytes and mesenchymal cells in both PAGA and cell type phylogeny analyses (Figure 6b-c), it suggests that evolutionary histories of these cell types are complex, potentially involving co-option of identity specification mechanisms from ancestral pinacocytes and mesohyl cells.

Among the *Sycon* choanocyte family of cells, choanoblasts map to *Spongilla* choanoblasts and cells identified as sperm cells in *Amphimedon*. This is not surprising, as sperm cells have been shown to be derived from choanocytes in many sponge species. Strikingly, *Sycon* choanocytes map to both choanocytes and archaeocytes (and archeocyte family cells, such as mesocytes) of *Amphimedon* and *Spongilla*. This finding supports the notion that in calcareous sponges choanocytes carry stem cell functions in calcareous sponges lacking archeocytes, and is in line with demonstrations of choanocytes and archaeocytes being able to directly differentiate into one another (Funayama 2010, 2013, 2018 and reviewed by Adamska 2021).

### Validating Haeckel’s Hypothesis: Cell Type Homology Between Sponges and Cnidarians

Finally, we mapped and compared *S. capricorn* cell types with cell types of two cnidarian model systems: freshwater hydrozoan, *Hydra vulgaris* (Siebert et al. 2019) and an anthozoan, a stony coral *Stylophora pistillata* (Levy et al. 2021). The vast evolutionary distance between these two species, as well as strikingly different levels of complexity between them increase our confidence in identified cell type homologies.

Overall, the mapping provided unequivocal support to Haeckel’s hypothesis of homology between sponge pinacoderm and cnidarian epidermis/ectoderm, as well as sponge choanoderm and cnidarian gastrodermis/endoderm (Figure 7b). The pinacoblasts mapped to epidermal cells in *Stylophora*, and epidermal stem cells in *Hydra*. Similarly, exo-pinacocytes mapped to epidermal cells and epidermal stem cells of *Hydra*, while also showing link with one of the coral gland cells, perhaps epidermal secretory cells. Interestingly, endo-pinacocytes mapped to *Stylophora* neurons. This is intriguing for two reasons – one, it might reflect deep evolutionary relationships between pinacocytes and ectoderm (which generally gives rise to neurons), but also because endo-pinacocytes (or epithelial cells lining the osculum) have been proposed to be sensory cells in sponges (Ludeman et al. 2014). However, as precursors of neurons and gland cells in *Hydra* showed similarity to some of choanocytes, as did *Hydra* cnidocytes (which are also a neuronal cell type), we are (not surprisingly) unable to provide insight into evolutionary origin of neurons using sponge dataset.

Overwhelmingly, the choanocyte family (including choanoblasts, choanocytes and apoptotic cells 2), mapped to gastrodermis family of cells. In *Stylophora*, these cells include gastrodermis itself, but also endodermal cells involved in algal symbiosis (the host cells). In *Hydra*, these include several types of gastrodermis and gastrodermis stem cells. Finally, choanoblasts, in addition to their link to *Stylophora* gastrodermis, show relationship with both coral and *Hydra* male germline, in a striking similarity to what we have discovered when comparing calcareous and demosponge cells.

Overall, our results are consistent with the last common ancestor of sponges and cnidarians (which might have been the same or different than the urmetazoan, depending on placement of ctenophores) being constructed from two epithelial cell layers, with the inner one involved in digestion and likely composed of collared cells. From this simple body plan, diversification of cell types and morphological complexity occurred in the diverse lineages of sponges, cnidarians and bilaterians, producing the spectacular diversity of animal forms we know today.

## Methods

### Collection of sponges

*S. capricorn* was collected by professional divers (*CrestDive*) at a depth of 3-5 meters in Jervis Bay, New South Wales, Australia. The sponges were then brought back to the laboratory facilities at ANU. All sponges were used in the first week after collection.

### High molecular weight DNA extraction and sequencing

High molecular weight genomic DNA (gDNA) was extracted from a single *S. capricorn* specimen using modified protocol after (Jones & Borevitz (2019), with the lysis performed in *Qiagen* RLT buffer and further fractionation using *Circulomix* Short Read Eliminator. The extracted gDNA fragment size ranged from 2,000 bp to 200,000 bp. gDNA was submitted to *Dovetail Genomics* for PacBio de novo sequencing. Dissociated cells of several additional *S. capricorn* individuals were fixed for 5min in 1.5% PFA were used for proximity ligation (Dovetail Hi-C protocol).

### Genome assembly and polishing

Dovetail Genomics performed scaffold-level genome assembly. To minimise errors in genome assembly and account for genetic variations between the RNA-Seq data and the reference genome, an initial genome refinement step was performed using de novo-assembled RNA-Seq transcripts. This transcriptome assembly was conducted with Trinity (Grabherr et al. 2011), followed by multiple rounds of genome correction using Pilon (Walker et al. 2014). The de novo assembled RNA-Seq transcripts were utilised solely for genome polishing. The polished genome was then used as a reference for RNA-Seq-guided prediction of gene and transcript structures, performed with StringTie (Pertea et al. 2015). Protein-coding regions were identified within the predicted transcripts using TransDecoder (Haas, Papanicolaou & Yassour 2024), and gene nomenclature along with functional annotation was assigned using Trinotate (Bryant et al. 2017). This annotation process incorporated GO terms and protein family classifications to provide comprehensive functional insights for each predicted gene.

### Living cell dissociation and cultivation

Each sponge individual was chopped by scalpels into tiny pieces and mechanically squeezed through 40 µm sieves (*pluriSelect*, 43-50040-01) in calcium and magnesium-free artificial seawater (Ca/Mg-FASW). Dissociated cells and spicules (heavier than cells) were separated by standing (2min), save the cell suspension and centrifuging (800g, 1min); repeat until no white spicule pellet was visible after centrifugation. The resuspended cells were transferred to 6-well plates, and 3 ml of artificial seawater (ASW) (*Aquaforest*, Reef Salt) contained 2 million cells per well. Cells were cultured at 16°C with replaced fresh ASW every 24 h. The cells and aggregates were observed and photographed every day using a *Leica* M205FA dissection microscope.

### Cell fixation and dissociation for single-cell sequencing preparation

Cells were fixed by a low concentration of methanal that combined with the lysis of acetic acid (modified from the ACME dissociation method (García-Castro et al. 2021)). Six sponge specimens were utilized in this study, with only the region encompassing the osculum and surrounding tissues, measuring approximately 1-1.5 mm, being sampled. The specimens were divided into two groups, with three sponges in each group (i.e., Group 1 and Group 2). Tissue samples from each group were collected and pooled at two distinct time intervals: 5-30 minutes and 65-90 minutes post-sampling, resulting in the formation of four replicates (i.e., Group1_5-30min, Group1_65-90min, Group2_5-30min, and Group2_65-90min). The sponge tissues, quickly minced in seawater, were placed in ACME with NaCl solution and rotated for 1 h, including two vigorous mechanical stirring and one 40 µm filtration. The cell suspension was allowed to stand on ice for 2min and collected supernatant, centrifuged (1000g, 5min, 4°C) and resuspended the pellet in 1 × PBS 0.5% BSA in the presence of RNAse inhibitor (40 U/mL) (*Roche*, 03335399001); repeated twice.

### Fluorescence-activated cell sorting (FACS) for single-cell sequencing preparation

The fixed cell suspension was stained by the nuclear dye DRAQ5 (*eBioscience*, 65-0880-92) (0.33 uL/mL). Cells were then sorted using a *BD* FACS Aria II (1.220) Cell Sorter at the Flow Cytometry Facility at ANU. Sorting was performed using the *BD* FACSDiva 8.0.3 Software, set up in 1-Way Purity mode, with a 100-μm nozzle. The gating of the FACS was decided to use forward scatter (FSC), side scatter (SSC), and DRAQ5 fluorescence (figure S2). The sorted cells have an average concentration of 107 cells/uL. The sorted cell suspension was stored on ice, and two small aliquots were removed for mixing with trypan blue (*Sigma*, T8154) to calculate the concentration or with the cytoplasmic dye Concanavalin-A conjugated with AlexaFluor 488 (Con-A 488) (*Invitrogen*, C11252) to confirm cell integrity under the Leica DM5500 compound fluorescence microscope.

### Single-cell RNA sequencing

After testing the cell concentration and confirming that the cells were intact, 43.2 uL cell suspension per replicate was immediately submitted to the Biomolecular Resource Facility at ANU. ‘Chromium Next GEM Single Cell 3ʹ GEM, Library & Gel Bead Kit v3.1 (4 rxns)’ (*10x Genomics*, PN-1000128) was used for the scRNA-sequencing library making of four cell replicates and sequenced using the Illumina NovaSeq6000 platform with SP-100 cycles.

### Pre-processing of 10x genomics scRNA-seq data

A custom reference made by the *S. capricorn* genome as well as gene annotation using the Cell Ranger (v7.1.0) ‘mkref’ pipeline (Zheng et al. 2017). Sample libraries were demultiplexed using the ‘mkfastq’ pipeline with the default setting and detected an average of 49,000 pairs of raw reads per loaded cell. Only exonic reads were aligned against the reference and transcript abundances were counted by the ‘count’ pipeline with the default setting. An average rate of 2886 cells per replicate was recovered.

### Pre-quality control

Downstream data from which cell doublets were removed by the default settings of the ‘ScDblFinder’ package were further processed by ‘Seurat (version 4.3.0)’ (Cao et al. 2022; Neavin et al. 2024). Gene features detected in less than 5 cells, as well as cells with less than 20 features, were removed by the ‘CreateSeuratObject’ function. Cells were then sorted by RNA quantity and number of genes expressed to remove outliers (figure S3a-c). This results in a final count matrix of 10,747 cells.

### De-batching and merging

After the four replicates of individual data were processed log normalization with default setting by the ‘NormalizeData’ function separately, the ‘FindVariableFeature’ function is used to figure out the top 2000 highly variable genes (HVGs) for anchor calculation, which uses the ‘FindIntegrationAnchors’ function. De-batching and merging of data were achieved by ‘IntegrateData’ algorithm.

### Dimension reduction

After screening out the top 2000 feature gene by ‘FindVariableFeature’ and ‘ScaleData’ functions, the ‘RunPCA’ algorithm was applied with default setting to perform Principal Component Analysis (PCA). ‘JackStraw’ and ‘ScoreJackStraw’ functions were then used to process the PCA scores with the top 20 dimensions selected.

### Clustering

’FindNeighbors’ and ‘FindClusters’ functions are used to calculate and identify cell clusters afterwards. Combining with ‘clustree’ algorithm (Zappia & Oshlack 2018), multiple top dimensions in ‘FindNeighbors’ and various resolutions in ‘FindClusters’ are set to visualize the stability of cluster splitting under different parameters. The top 16 dimensions and 0.12 resolution were finally determined as the best parameters (figure S4a-b). Cluster ‘2’ is further isolated and re-clustered into two clusters (figure S4c). A total of 11 clusters were initially detected. ‘FindAllMarkers’ function was further applied to identify the marker genes of each cell type, which was used to identify and annotate cell types. After confirming the cell type identity, the clusters were re-named by ‘RenameIdents’ function and visualized with the UMAP by ‘RunUMAP’ function.

### Post-quality control

The clustering of four replicates was compared on Umap based on gene expression. The results showed a high degree of overlap between the replicates (figure S3d-e). To further compare the cell ratio of four independent replicates without assuming a normal distribution, a Kruskal-Wallis test was performed to determine whether there are significant differences between replicates either across the atlas dataset or within a specific cell type family. The results showed that there was no significant difference in the cell ratio between replicates (figure S5).

### RNA probes designing and synthesis

The RNA transcript sequence of cell-type marker genes (figure S6) was used to design 500-1000bp antisense probes. T3-polymerase recognition sequence was added upstream of the 3’ end of the fragments. Templates were ordered from *Integrated DNA Technologies* (*IDT*) as gBlocks Gene Fragments (Appendix III). The ordered templates were mixed with DIG DNA Labeling Mix (*Roche*, 11277073910), RNAase inhibitor, T3 RNA polymerase and related buffer (*Roche*, 11031163001) and incubated at 37°C for two hours. The products were purified using the RNeasy MinElute Cleanup kit (*QIAGEN*, 74204).

### In situ hybridization

ISH was carried according to Fortunato et al. (2012): the sponge tissues including oscula were fixed with 4% paraformaldehyde/0.05% glutaraldehyde in MOPS buffer (MOPS fix) for 1 h at room temperature (RT) and stepped into 70% ethanol. Following storage at −20°C, samples were rehydrated and incubated in 7.5 µg/mL of proteinase K (*Sigma*, 1.24568) at 37°C for 10 minutes and the reaction terminated with glycine. Tissues were acetylated by 1.5-3uL/mL acetic anhydride (*Sigma*, 320102) in 0.1M Triethanolamine (*Thermo Scientific*, J63793.AK). After being fixed again with MOPS fix and washed, the tissues were incubated with hybridisation buffer (HB) at 51°C for 1.5-3 h for pre-hybridisation. RNA probes were mixed in HB at the designed concentration (Appendix III) and preheated at 70°C for 10 minutes. The HB in which the tissues were incubated was replaced by the HB with probes and incubated at 51°C for another 12-18 h. To increase the stringency of the probe to target binding, tissues were incubated with formamide (*Sigma*, 47671) and saline-sodium citrate (*Sigma*, S6639), which were used in high to low concentrations with Tween-20 at 51°C. After a series of washes, tissues were incubated in 2% blocking buffer in maleic acid buffer (MAB) for 1 h at RT to block nonspecific binding sites. Anti-Digoxigenin-AP (*Roche*, 11093274910) at 1:5000 was then added and incubated overnight at 4°C. After several washes to remove the antibody for 3 hours, the tissues were washed with Mg-free AP buffer. Tissues were then immersed with BM-Purple (*Roche*, 11442074001) at RT and kept in the dark to allow the probe-bound areas to develop colour (purple). The stained tissues were observed and photographed using the *Leica* M205FA dissection microscope.

### Sectioning

In order to investigate the characteristics of the stained cells more clearly, including their specific location and shape, the stained tissues were sectioned into slides and further observed. After a 60-minute incubation in 175 mM EDTA and 175 mM EGTA in PBS and washes to remove spicules, tissues were immersed in 10% sucrose in TBST (Tris-buffered saline with 0.1% Tween-20 detergent) for 30 minutes to enhance sectioning performance. Place the tissue without droplets into a 1cm * 1cm * 0.5cm Tissue-Tek Cryomold (*SAKURA*, 4565) and fill it with Tissue-Tek O.C.T. (optimal cutting temperature) (*SAKURA*, 4583). After the cryomold with tissue was completely frozen, the square tissue block was taken out and mounted on the stage of a *Leica* CM1860 cryostat microtome set at −30°C. The tissue block was sectioned into slices of 15-30 μm thickness, and cryofilms (type IVD(16UF), *SECTION-LAB, Co., Ltd.*) were used as a carrier to protect the sections’ integrity (Kawamoto & Kawamoto 2021). The cryofilms with sections attached were washed and then embedded in glycerol. The cryofilms were then made into slides, observed and recorded using DIC (differential interference contrast) under the *Leica* DM5500 compound fluorescence microscope.

### Gene ontology enrichment analysis

Gene ontology (GO) enrichment analysis was performed to predict the functions of cell types based on their marker genes. Among the top 50 marker genes in each cell type, genes with high statistical significance and high fold change were retained (adjust p-value < 0.001 & average of log2FC > 1). Only genes that can be matched with the GO term under the consideration of protein family were retained. Gene set variation analysis (GSVA) was used to estimate whether these marker gene lists were significantly enriched in their corresponding cell types (figure S7) (Haenzelmann, Castelo & Guinney 2013). The unannotated genes from the filtered top genes were also analysed using GSVA for each cell type, yielding similar expression patterns to the annotated genes (figure S8). This consistency demonstrates that the annotated genes accurately reflect the true marker gene expression patterns. For each cell type and its retained annotated marker genes, GO-seq was used to account for the gene length bias in the detection of over-representation to judge the significance of each GO term (Young et al. 2010). Only GO-term annotations with p-value ≤ 0.05 are retained.

### Transcription factor analysis

All genes annotated with the GO term GO:0003700, DNA-binding transcription factors, were considered transcription factors of *S. capricorn*. In addition, Steinegger & Soeding (2017)’s ‘mmseqs easy-rbh’ tool that conducts reciprocal best hit (RBH) was used to identify orthologous genes of transcription factors and signalling molecules between *S. capricorn* and its congeneric species, *S. ciliatum*. The published protein sequences of transcription factors and signalling molecules of *S. ciliatum* were used to hit all the protein sequences of *S. capricorn* (Adamska et al. 2007; Fortunato et al. 2014; Leininger et al. 2014; Fortunato, Adamski & Adamska 2015; Fortunato et al. 2016). Even though some genes of *S. capricorn* are annotated with the same gene name in other species, they are still distinguished by a number as a suffix to avoid confusion. The expression of genes of signalling molecules and transcription factors is demonstrated in the form of dot plots (figure S16).

### RNA velocity analysis

To predict the temporal dynamics of single cells based on their gene expression, the ‘velocyto’ pipeline was used to describe RNA splicing dynamics (La Manno et al. 2018). Alignment results generated by Cell Ranger were filtered based on the retained cell barcodes using the ‘subset-bam’ pipeline made by 10X Genomics and converted to Loom matrix using the ‘velocyto run’ pipeline. The parameters of the ‘RunVelocity’ function were set to deltaT = 1, kCells = 15, and fit.quantile = 0.2, and RNA velocity was finally visualised on UMAP by the ‘velocyto.R (v.0.6)’ package.

### Partition-based graph abstraction (PAGA) connectivity graph

The strength of connections between individual cells was calculated and visualized per cell type. Based on the PAGA algorithm proposed by Wolf et al. (2019), the similarity of each pair of cells was calculated and connected to form a network with different connectivity scores. This PAGA network was conducted and simplified using Python models built with the ‘SCP’ package (Zhang 2023). The threshold of connectivity score is set to 0.03; values below 0.1 are displayed in grey, and values above 0.1 are displayed in black. The thickness of the connection line is used to distinguish the strength of the connection further.

### Cell Type Phylogeny

In order to intuitively predict and display the relationship between cell types, a neighbour-joining tree, which was modified from Musser et al. (2021)’s methods based on bootstrapping and the high variant genes (HVGs) of each cell type, was estimated. The estimation bootstrapping was repeated 10,000 times and showed the percentage of repetition frequency, which was the robust rate of the branch, using the R package ‘ape’ and ‘phytools’ (Paradis & Schliep 2019; Revell 2024). The trees’ branching structures were similar (i.e., robust clade structure) when using the average expression of the top 30% - 100% of the entire HVGs. The published tree estimation with the top 1280 HVGs was based on the highest repeatability of nodes near the centre of the tree.

### CytoTRACE

Developmental potential can be deduced by the number of detectably expressed genes per cell. The higher the transcriptional diversity, the less differentiation there is usually. The extracted raw gene expression matrix of each scRNA-seq sample was input into the ‘iCytoTRACE’ function of the ‘CytoTRACE’ package in parallel (Gulati et al. 2020). Each cell’s degree of deducted differentiation was scored as a value from 0 to 1 and visualized using ‘ggplot2’ (Hadley 2016).

### Self-assembling Manifold Mapping (SAMap)

To compare single-cell datasets from sponges and cnidarians, a specialized method known as Self-Assembling Manifold mapping (SAMap) (Tarashansky et al. 2021) was utilized. Specifically, the 10X *S. capricorn* dataset (10,747 cells) generated in this study, the 10X *Spongilla lacturis* dataset (10,106 cells) (Musser et al. 2021), the MARS-seq adult *Amphimedon queenslandica* dataset (3,870 cells) (Sebé-Pedrós et al. 2018), the Drop-seq *Hydra vulgaris* dataset (20,789 cells) (Siebert et al. 2019; Levy et al. 2021), and the MARS-seq *Stylophora pistillata* dataset (16,080 cells) (Levy et al. 2021) were aligned. To execute SAMap, raw count data from each dataset and a comprehensive list of homologous gene pairs were required. Homologous gene pairs were identified using reciprocal tblastx analysis with an e-value threshold of 1e^-6^, implemented through the BLAST mapping script provided within the SAMap framework. Subsequently, a graph-stitching procedure implemented in SAMap was applied to link the datasets and refine the list of homologous gene pairs by retaining those exhibiting overlapping expression patterns across the aligned manifold. This refinement process was repeated for a defined number of iterations until convergence of the aligned manifold was achieved. SAMap was executed with the following settings: NH1 = 2, NH2 = 2, and NUMITERS = 50. The NUMITERS parameter, which defines the number of iterations performed, was increased from the default value of 3 to 50 to ensure sufficient convergence of the aligned manifold. The parameters NH1 and NH2 specify the number of steps away from each cell in datasets 1 and 2, respectively, to be included in its neighborhood. Values of NH1 = 2 and NH2 = 2 were used instead of the default value of 3, consistent with the implementation of Musser et al. (2021). In Musser et al. (2021)’s study, neighborhood sizes were reduced based on recommendations from the SAMap tutorial vignette to prevent overlap between distinct cell-type neighborhoods in smaller datasets, and the same approach was followed here. Sankey plots were generated using the Sankey_plot function implemented in SAMap, and the gene pairs driving the connections displayed in the Sankey plots were extracted using the GenePairFinder function within SAMap.

## Author contributions

Study design: D.P., D.R., M. Adamska; Methodology development: D.P., D.R.,C.C, R.R, M. Adamski, M. Adamska; Laboratory experiments: D.P., M. Adamska, Bioinformatic analyses: D.P., D.R., M. Adamski; Visualization: D.P., D.R.; Supervision: M. Adamski, M. Adamska; Writing: D.P., D.R., M. Adamska.

## Acknowledgments

We are grateful to the following ANU facilities for their technical support for this project: Ecogenomics and Bioinformatics Lab (EBL), Flow Cytometry Facility (FCF), Biomolecular Resource Facility (BRF), and RSB IT Service Team. We greatly appreciate Sue Newson (*CrestDive*) for sponge sampling; Harpreet Volhra for advice and assistance with FACS, Giel van Noorden for cryotome sectioning advice; Yitang Du for help with Figure 1c, Farid Rahimi for microscopy and centrifuge support; Tiannan Gao for field work assistance. M. Adamska deeply appreciates advice given by Niccy Aitken and Ashley Jones in development of gDNA isolation protocol, Nathan Kenny in selection of the single cell fixation protocol and Noriko Funayama for stimulating discussions on sponge cell biology. The black silhouettes in the phylogenetic tree (Figure 1a) were sourced from *PhyloPic*, and the tree was generated using *BioRender*. All graphical elements are reproduced under the appropriate licenses.

