## Supplementary figures for "Calcareous sponge cell atlas provides support to homology between sponge and eumetazoan body plans"

**Appendix I (supplementary figures)**

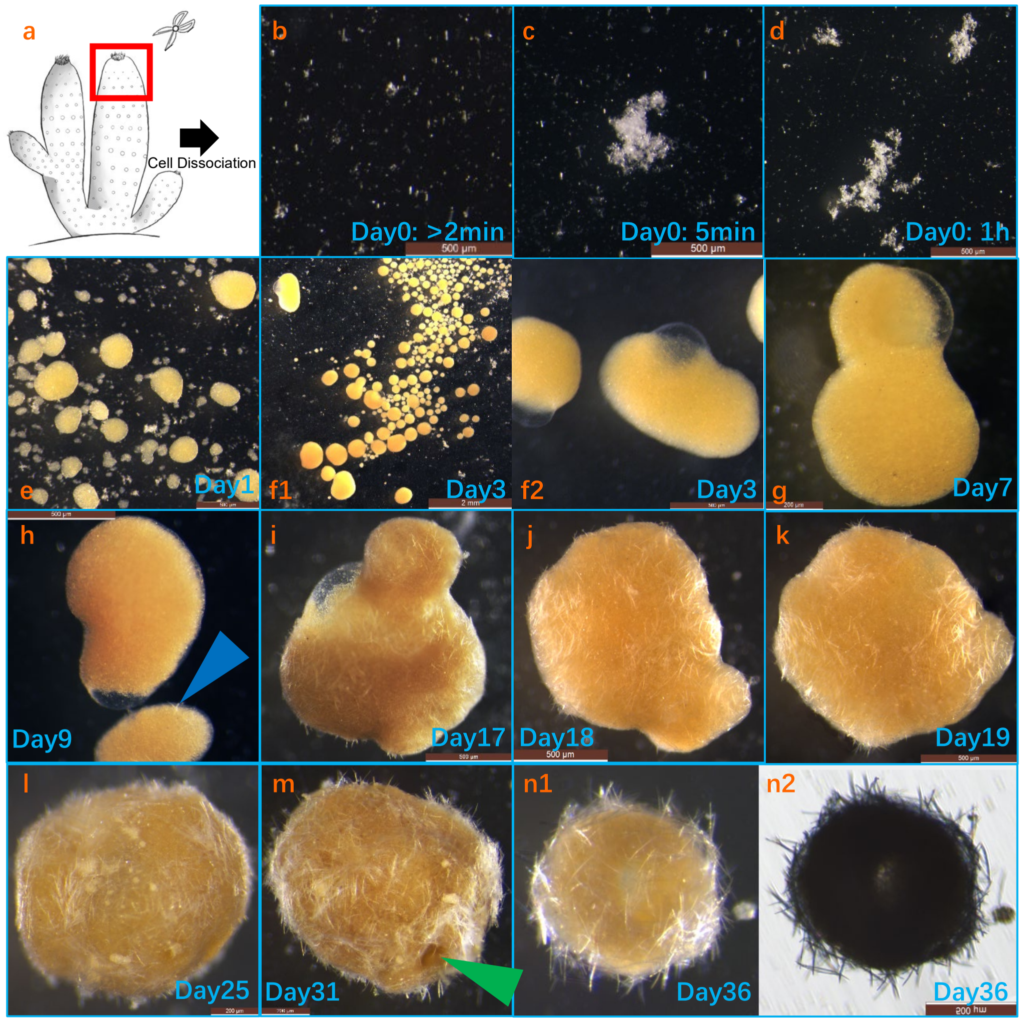


**figure S1. Reaggregation from the dissociated single cells to a juvenile in *S.capricorn*.** **a**, Sponge fragmentation and cell dissociation schematic diagram. **b-e**, Reaggregation is activated from the moment of dissociation and gradually forms yellow multicellular aggregates. **f1-g**, Clear bubbles appear on the surface of multicellular aggregates from day 3, which are called lacunae. Multicellular aggregates continue to aggregate with each other, whether or not they have lacunae. **h**, Multicellular aggregates develop and generate spicules (as indicated by the blue arrow). **i-l**, Spicules are abundantly spread over the surface of the aggregates. The initially translucent, membrane-like surface of the lacunae gradually thickens over time, eventually becoming fully enclosed within the interior of the aggregate and positioned centrally. As this process progresses, the aggregate gradually re-establishes a rounded morphology, culminating in the formation of a hollow spheroid. **m**, Osculum-like structures (as indicated by the green arrow) appeared on the surface of the aggregates, connecting the inner lacunae with the outside. **n1-n2**, From day 36, the aggregate appears as a hollow cavity, which is the juvenile-like stage of *Sycon*. **n1** (front-illumination) and **n2** (back-illumination) are showing the same juvenile-like aggregate. Notably, the larger (upper) aggregate observed in panel **h** represents the same aggregate in panel **g**. In panel **i**, the two aggregates have merged to one from panel **h**, and at all subsequent timepoints (**i-n**) the panels track this single aggregate, which does not incorporate any additional aggregates thereafter.

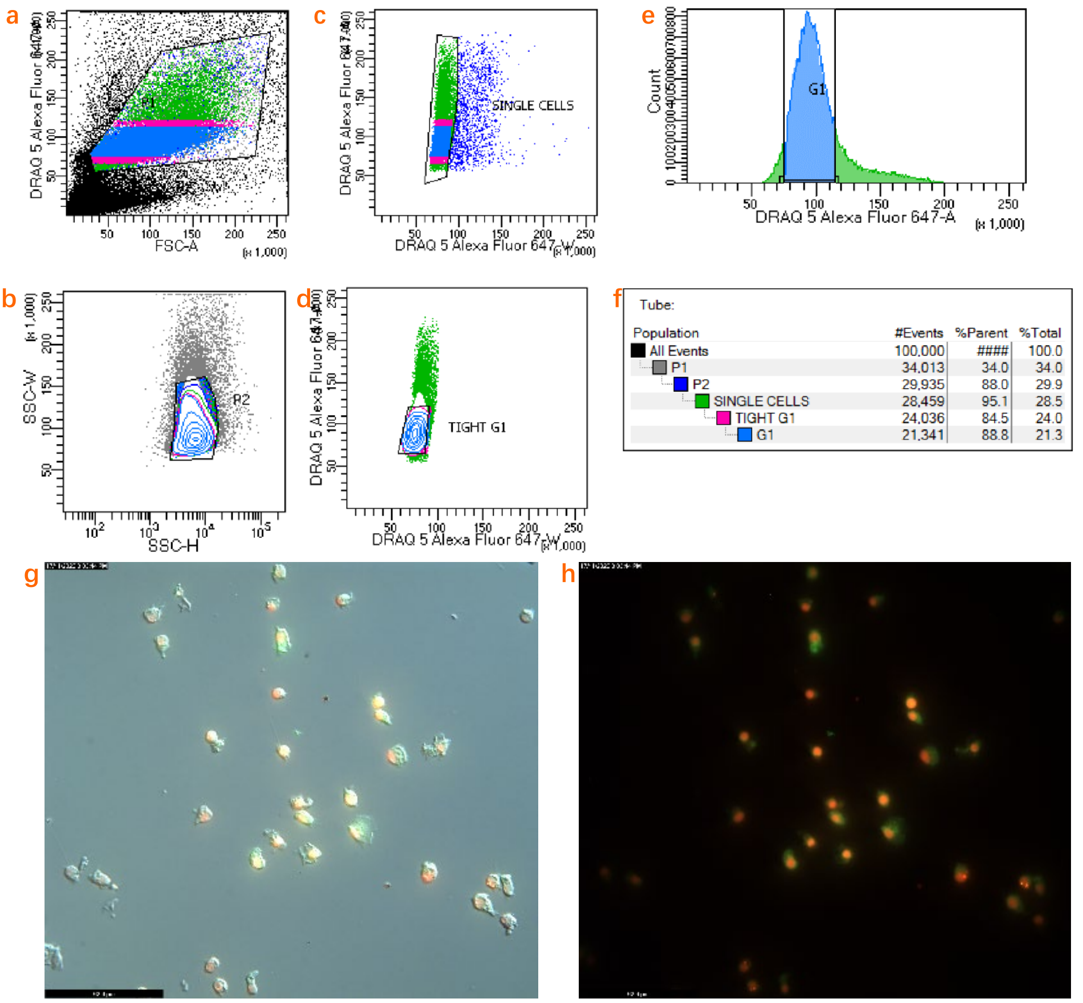


**figure S2. FACS gating for single cells and the sorted single cells with DRAQ5 and Con-A 488 fluorescence. a-e**, Gate settings were used to avoid the collection of (**a-b**) non-cell noise, broken cells, (**c**) cell doublets or clusters, and (**d-e**) cycling cells with duplicated DNA; **f**, The structure of the gate population; **g,** Combined DIC and fluorescence imaging of sorted cells; **h,** Fluorescence imaging only of sorted cells. Various cell types, especially flat pinacocytes and flagella-bearing choanocytes, are easily distinguished. Red fluorescence is nuclear dye DRAQ5; green fluorescence is cytoplasmic dye Con-A 488.

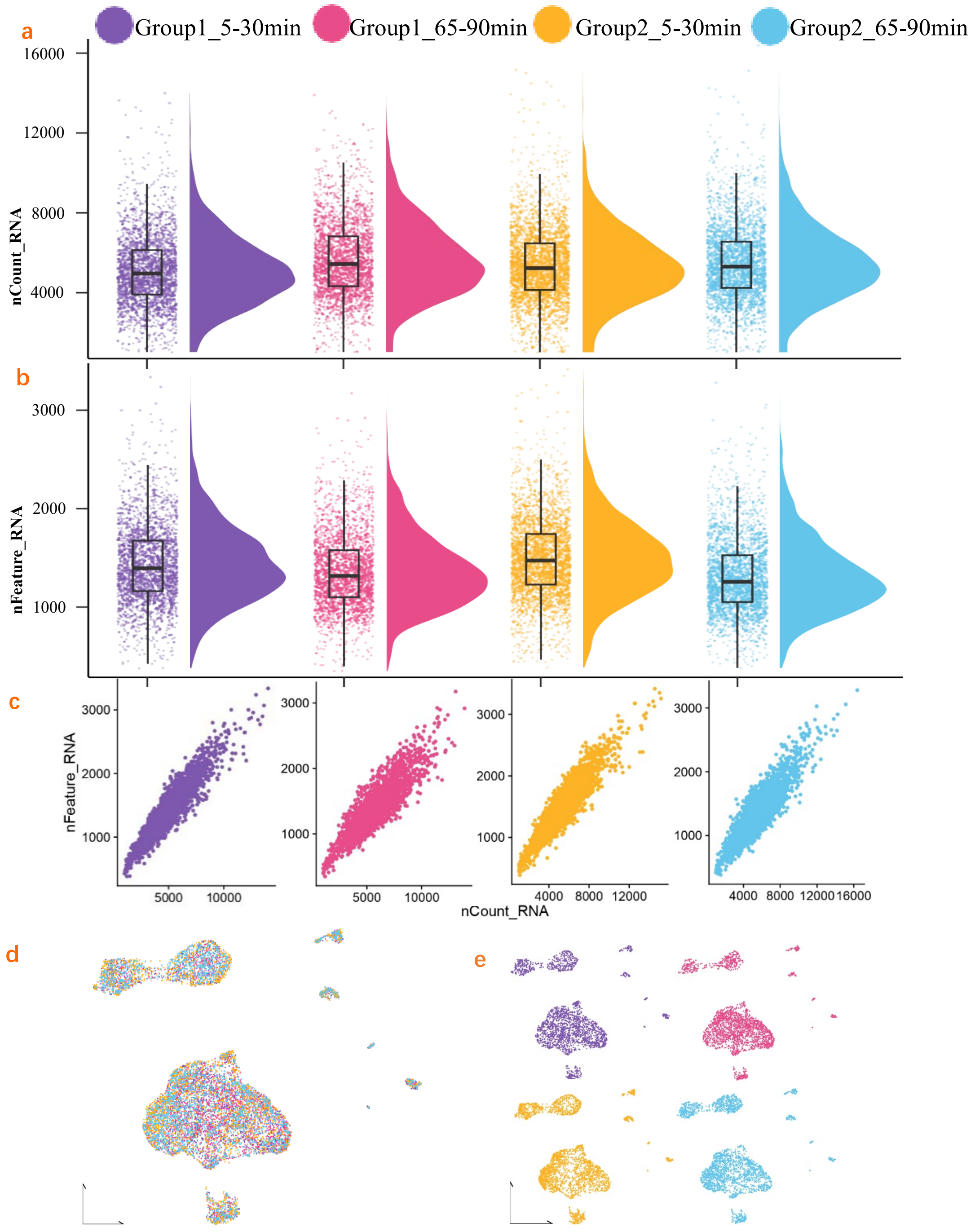


**figure S3. The reproducibility of the four scRNA-seq replicate groups.** The RNA quantity (**a**, 'nCount_RNA'), number of expressed genes (**b**, 'nFeature_RNA'), and both parameters (**c**) of each cell per group after quality control are shown. The distribution of the cells of four groups after merging (**d**) and splitting (**e**) on UMAP. Each color corresponds to a group.

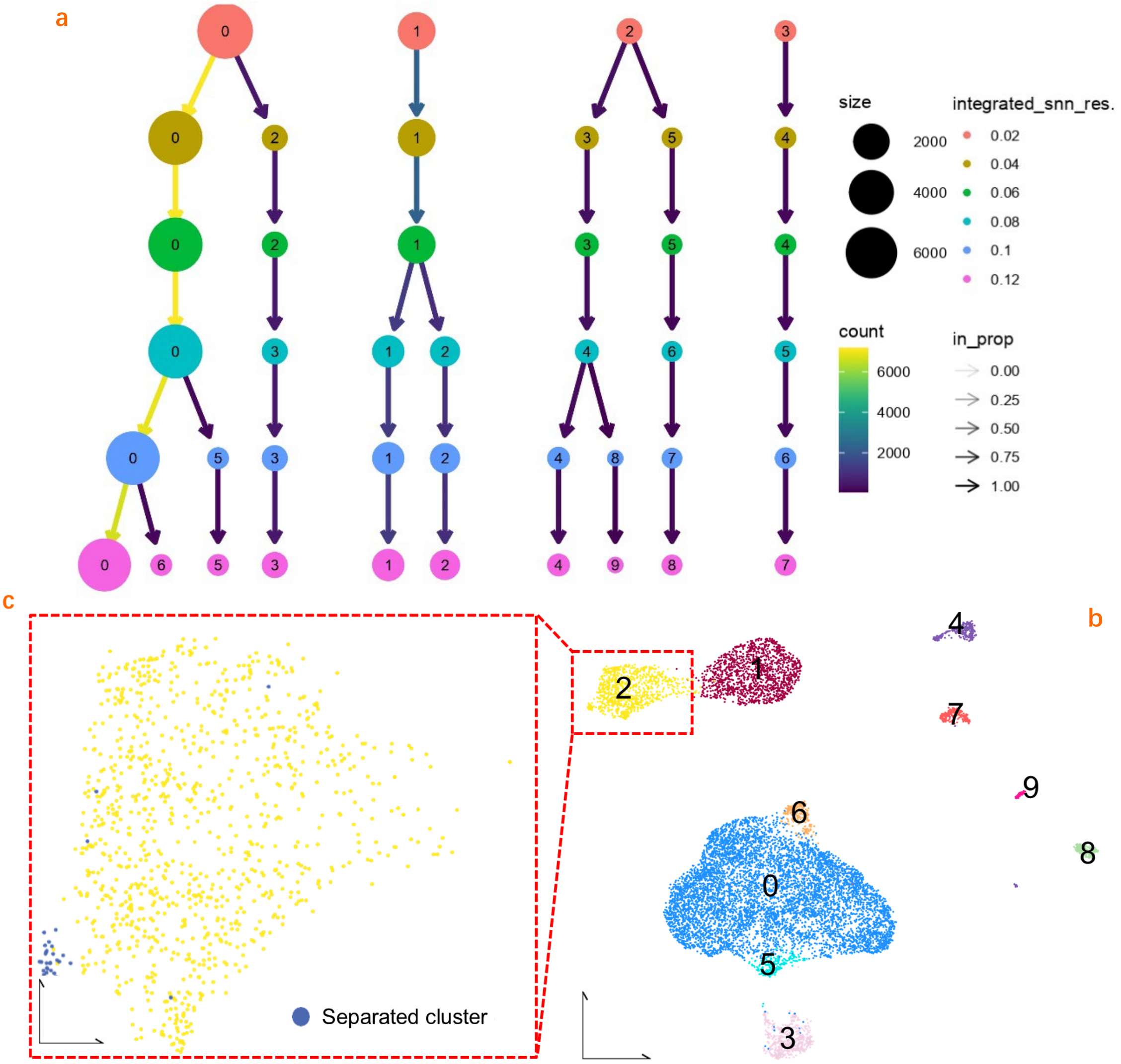


**figure S4. Cell cluster detection and secondary splitting. a**, The cluster tree illustrates how cells were progressively and stably partitioned into ten major clusters (labelled 0-9) as the resolution increased during graph-based clustering. Node size as well as arrow colour reflect the number of cells in each cluster, while node colour represents the corresponding clustering resolution from 0.02 to 0.12; **b**, UMAP projection showing the spatial distribution of the ten main clusters (bottom right); **c**, One of these clusters, cluster 2 (yellow), was further computationally subdivided into two subclusters (bottom left). Each colour in the UMAP corresponds to a distinct cell cluster, totalling eleven clusters after secondary splitting.

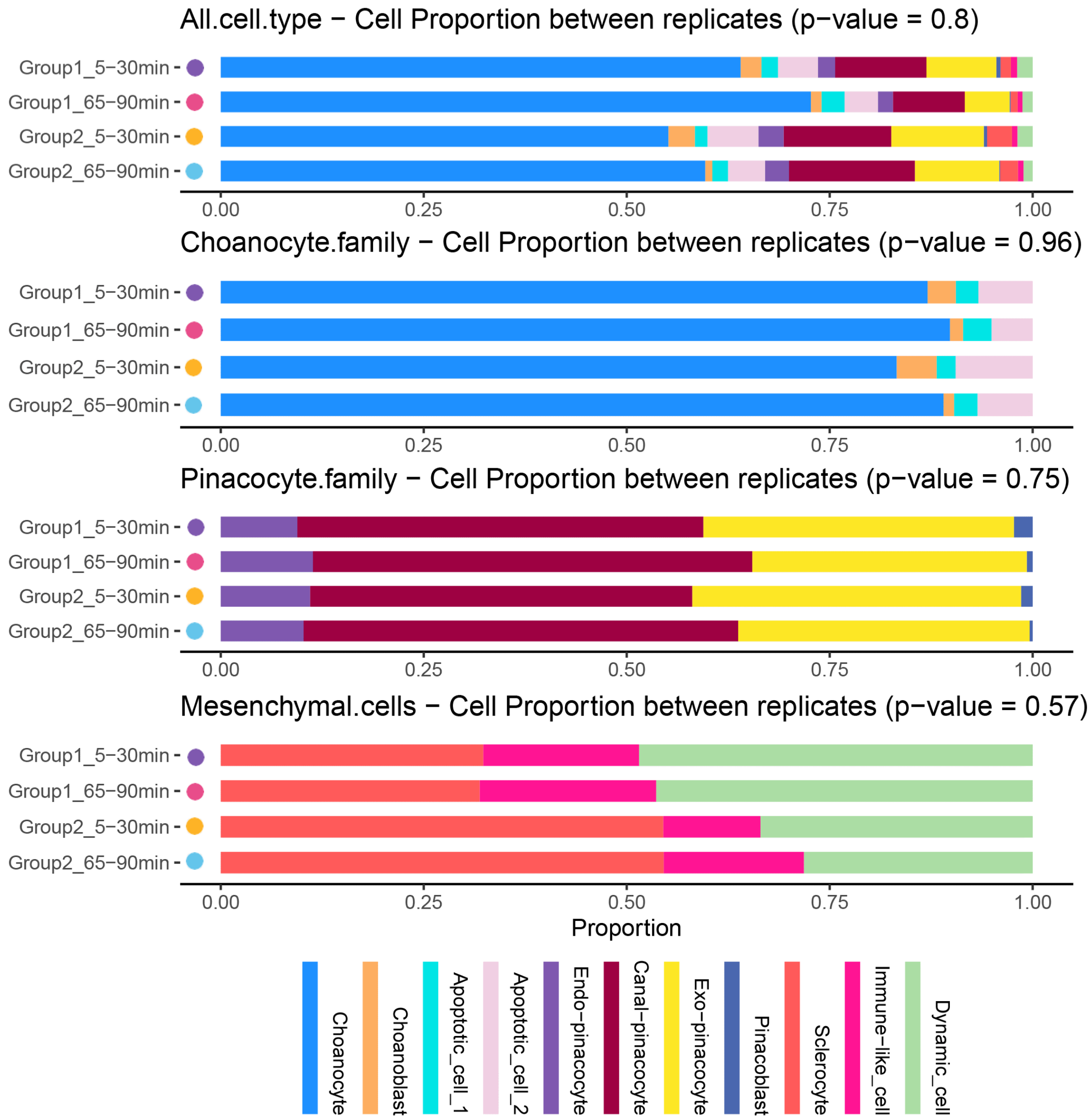


**figure S5. Comparing the cell ratio of four independent replicates either across the atlas dataset or within a specific cell type family.** Kruskal-Wallis tests were performed to determine whether there were significant differences among the replicates. The p-value has been annotated in the title of each bar graph, and they are all greater than 0.05. This indicates that there is no significant difference among the four replicates.

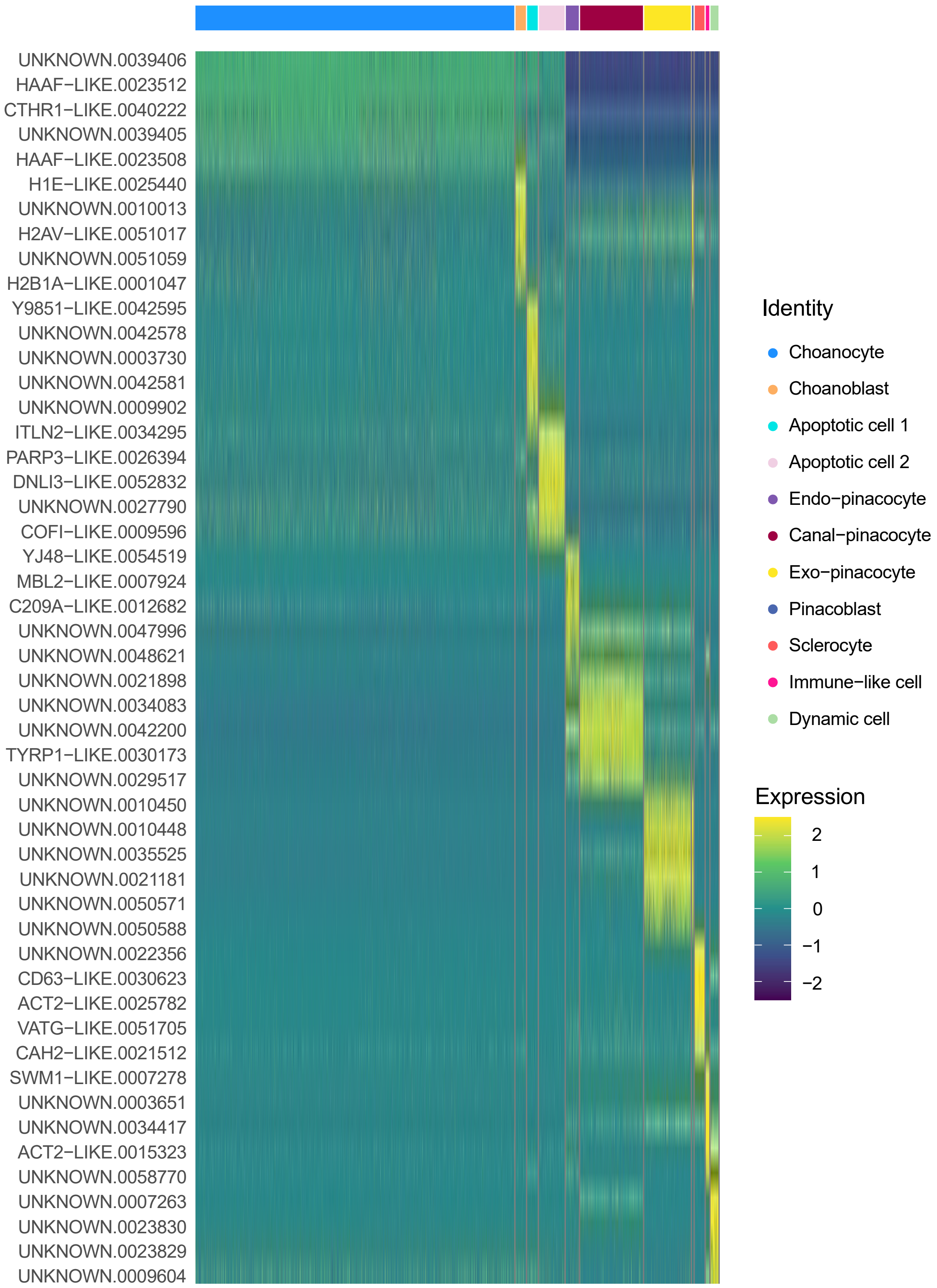


**figure S6. Expression heatmap of top 5 marker genes of each cell type.** The rows of the heat map represent genes, and the columns represent cells. Yellow indicates that the gene is highly expressed in the corresponding cells.

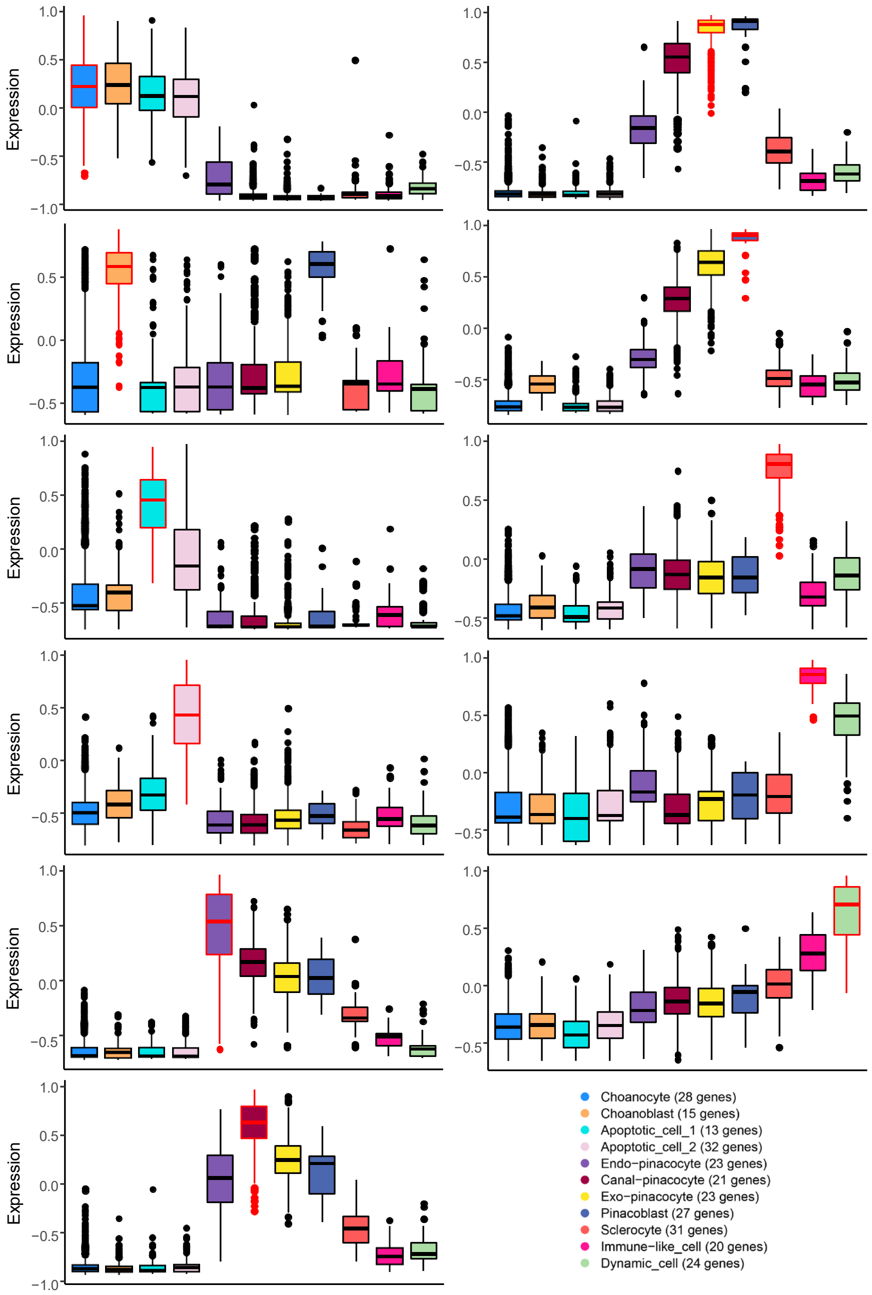


**figure S7. GSVA evaluates the specificity of marker gene lists of each cell type that are used for GO analysis in Figure 3.** Each plot compares the expression (i.e., GSVA score) of the marker gene list of the cell type highlighted by the red border in each cell type. The corresponding genes used in each plot are the genes annotated by GO terms among the top no more than 50 genes of the highlighted cell type. Some genes were screened out by quality control from the top 50 genes. The final number of genes used is reflected in the legend in the lower right corner, and the lists of gene IDs are presented in Appendix IV. The higher the GSVA score, the more specific the gene list is expressed in the corresponding cell type.

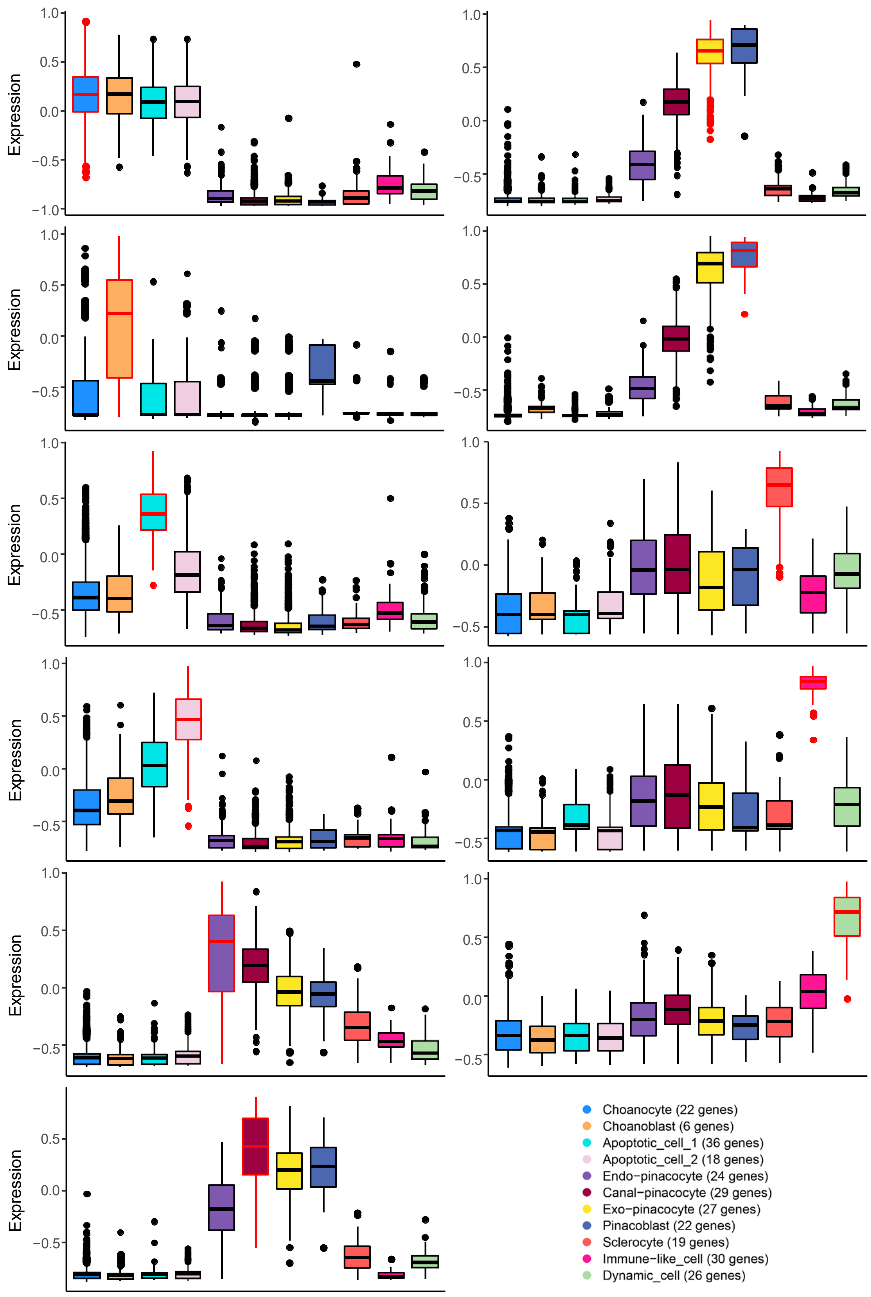


**figure S8. GSVA evaluates the specificity of marker gene lists of each cell type that cannot be annotated by GO terms among the top genes.** The gene lists used here are complementary to the ones used in figure S8. Each plot compares the expression (i.e., GSVA score) of the marker gene list of the cell type highlighted by the red border in each cell type. The corresponding genes used in each plot are the genes that cannot be annotated by GO terms among the top no more than 50 genes of the highlighted cell type. Some genes were screened out by quality control from the top 50 genes. The final number of genes used is reflected in the legend in the lower right corner, and the lists of gene IDs are presented in Appendix IV. The higher the GSVA score, the more specific the gene list is expressed in the corresponding cell type.

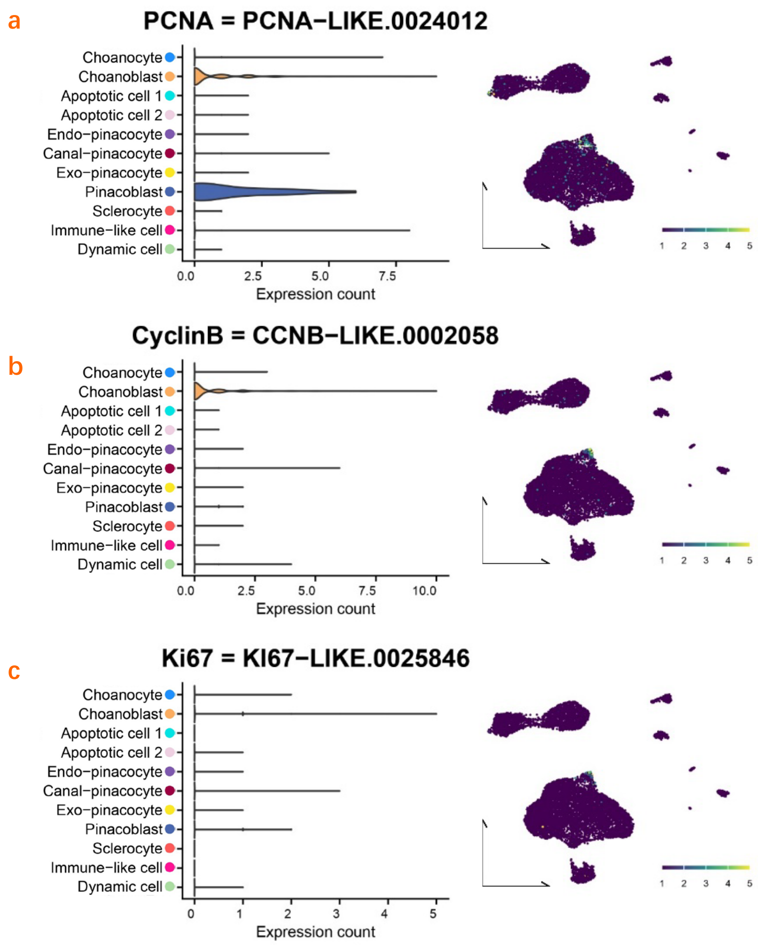


**figure S9. Genes related to multiple cell cycle phases were highly expressed in blast cells (choanoblast and pinacoblast) partially.** The average transcript counts of *PCNA* (**a**), *CyclinB* (**b**), and *Ki67* (**c**) in each cell type are shown in the left violin plots. Cells expressing the indicated genes are visualized in the right UMAP. Yellow indicates that the gene is highly expressed in the corresponding cells.

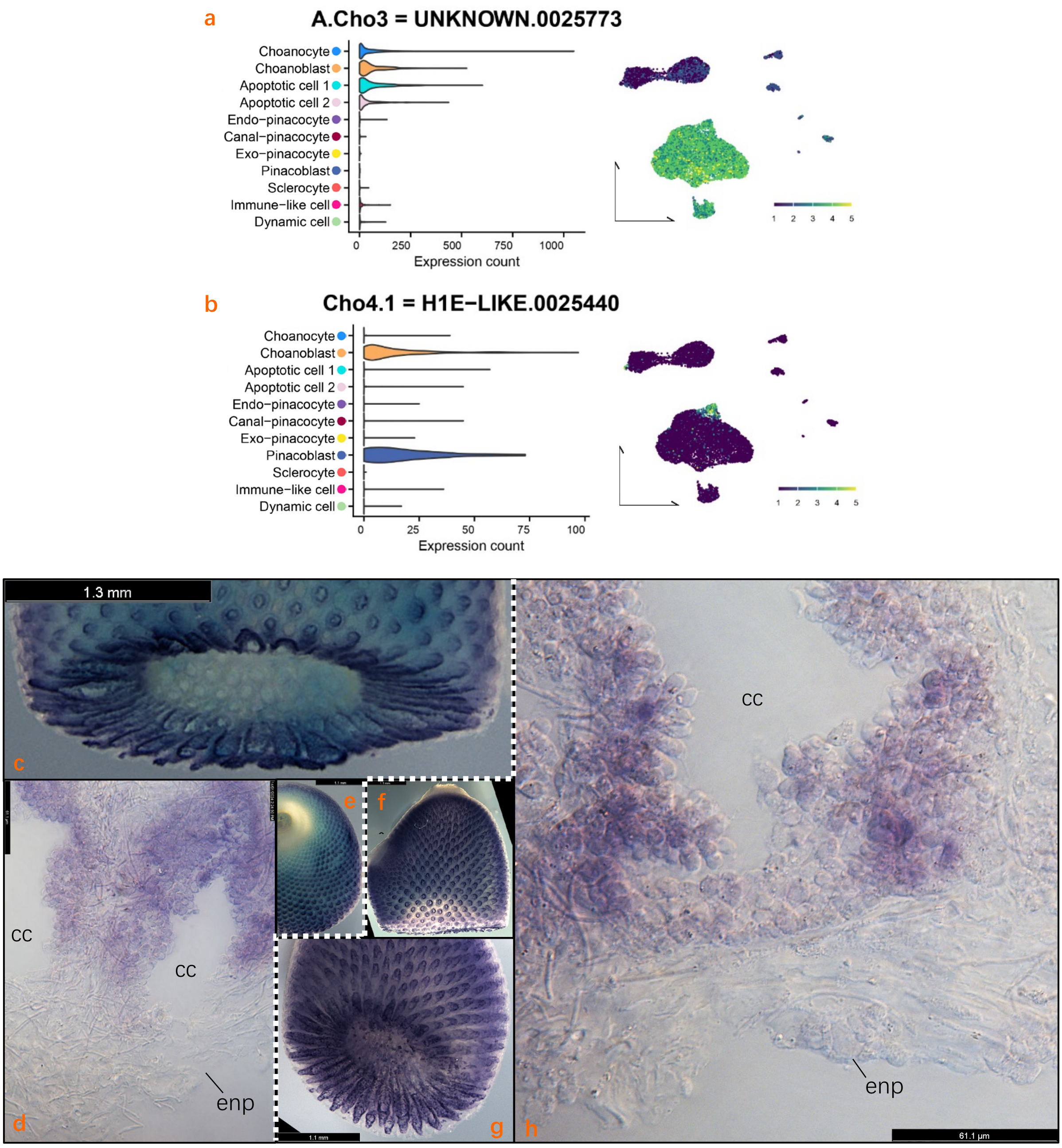


**figure S10. Gene expression visualization of the marker genes of the choanocyte family and proliferative cells. a**, **c-e**, The expression level and cell distribution of the marker gene of the choanocyte family; **b**, **f-h**, The expression level and cell distribution of the marker gene of the proliferative cells. The purple colour in the hybridization results indicates that the cells at that location highly express the specified marker gene.

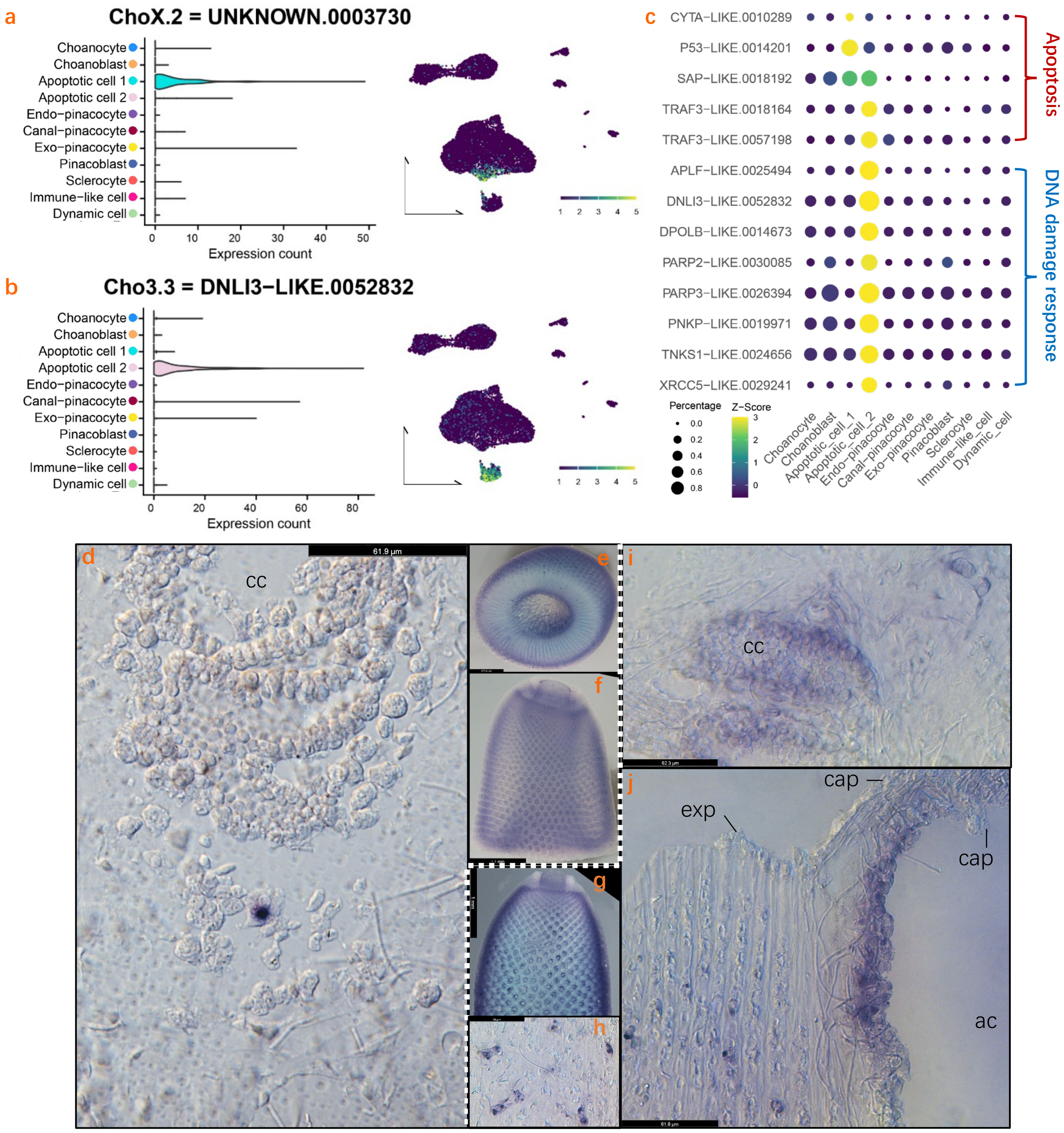


**figure S11. Gene expression visualization of the marker genes of apoptotic cells. a**, **d-f**, The expression level and cell distribution of the marker gene of apoptotic cell 1; **b**, **g-j**, The expression level and cell distribution of the marker gene of apoptotic cell 2. The purple colour in the hybridization results indicates that the cells at that location highly express the specified marker gene; **c**, The expression ratio among cell types (Z-score) and within each type (Percentage) of the designated marker genes of apoptotic cell 1 and apoptotic cell 2. Gene functions are categorised on the right. There are a large number of unknown genes that cannot be annotated in the top marker genes of apoptotic cell 1. Therefore, only few genes are shown here.

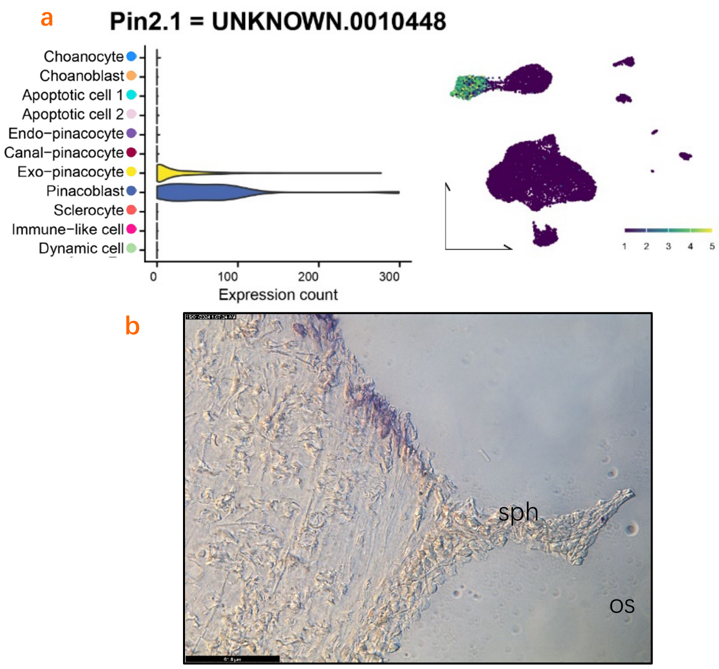


**figure S12. Gene expression visualization of the marker gene of exo-pinacocytes and pinacoblasts. a-b**, The expression level and cell distribution of the marker gene of exo-pinacocytes and pinacoblasts. Exo-pinacocytes do not have a unique marker gene independent of pinacoblasts. The purple colour in the hybridization results indicates that the cells at that location highly express the specified marker gene.

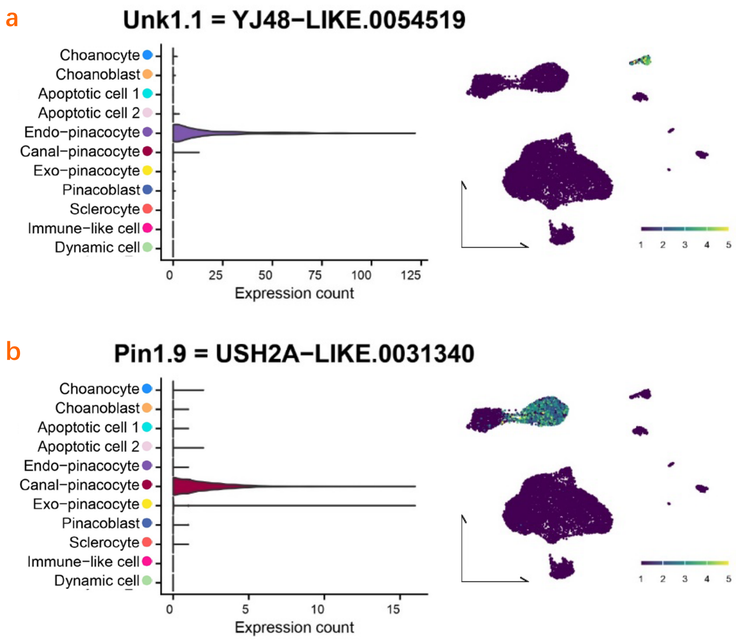


**figure S13. Gene expression visualization of the marker genes of endo-pinacocytes and canal-pinacocytes. a**, The expression level and UMAP cell distribution of the marker gene of endo-pinacocytes; **b**, The expression level and UMAP cell distribution of the marker gene on UMAP of canal-pinacocytes.

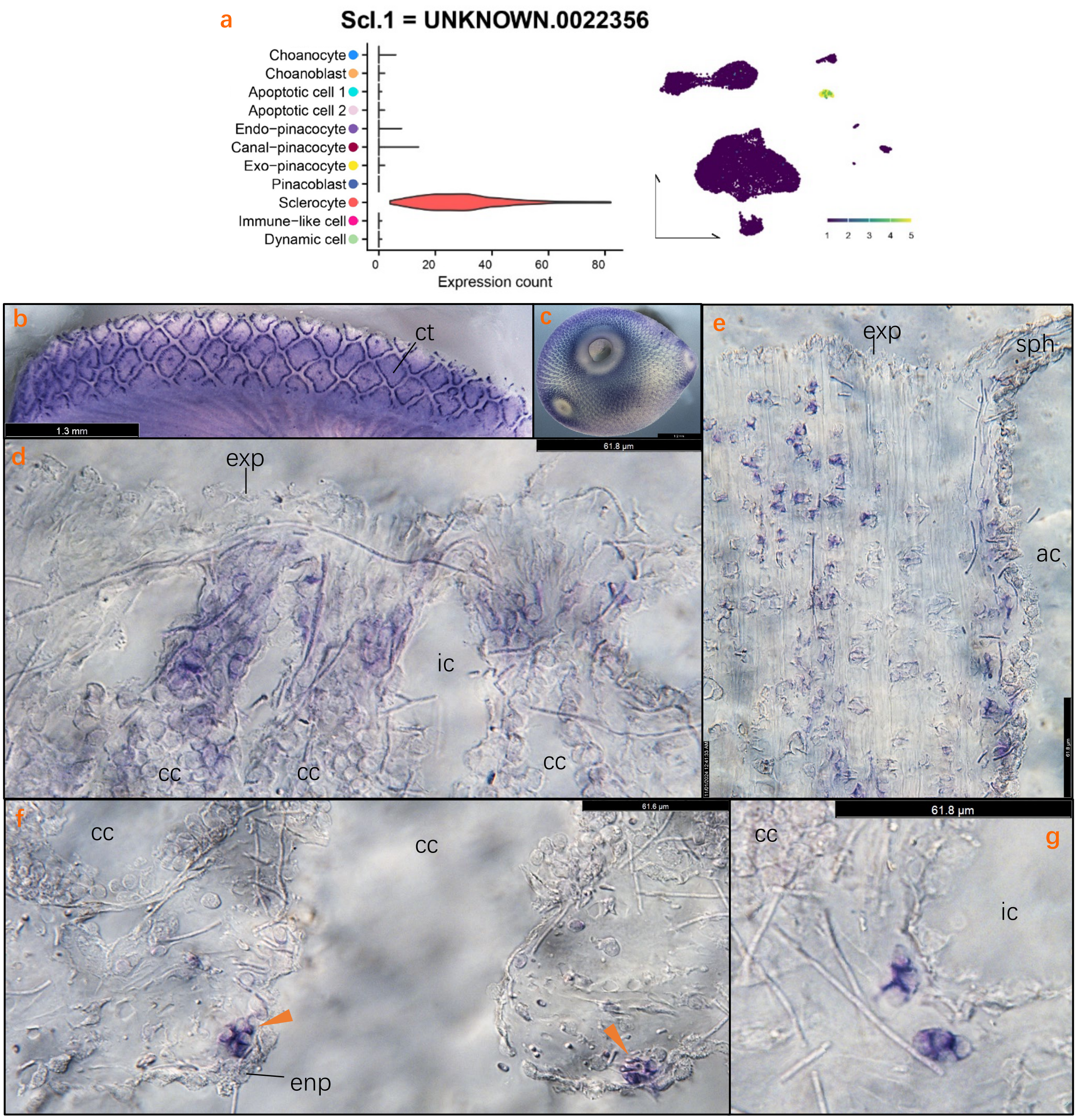


**figure S14. Gene expression visualization of the marker gene of sclerocytes. a-g,** The expression level and cell distribution of the marker gene of sclerocytes. The purple colour in the hybridization results indicates that the cells at that location highly express the specified marker gene.

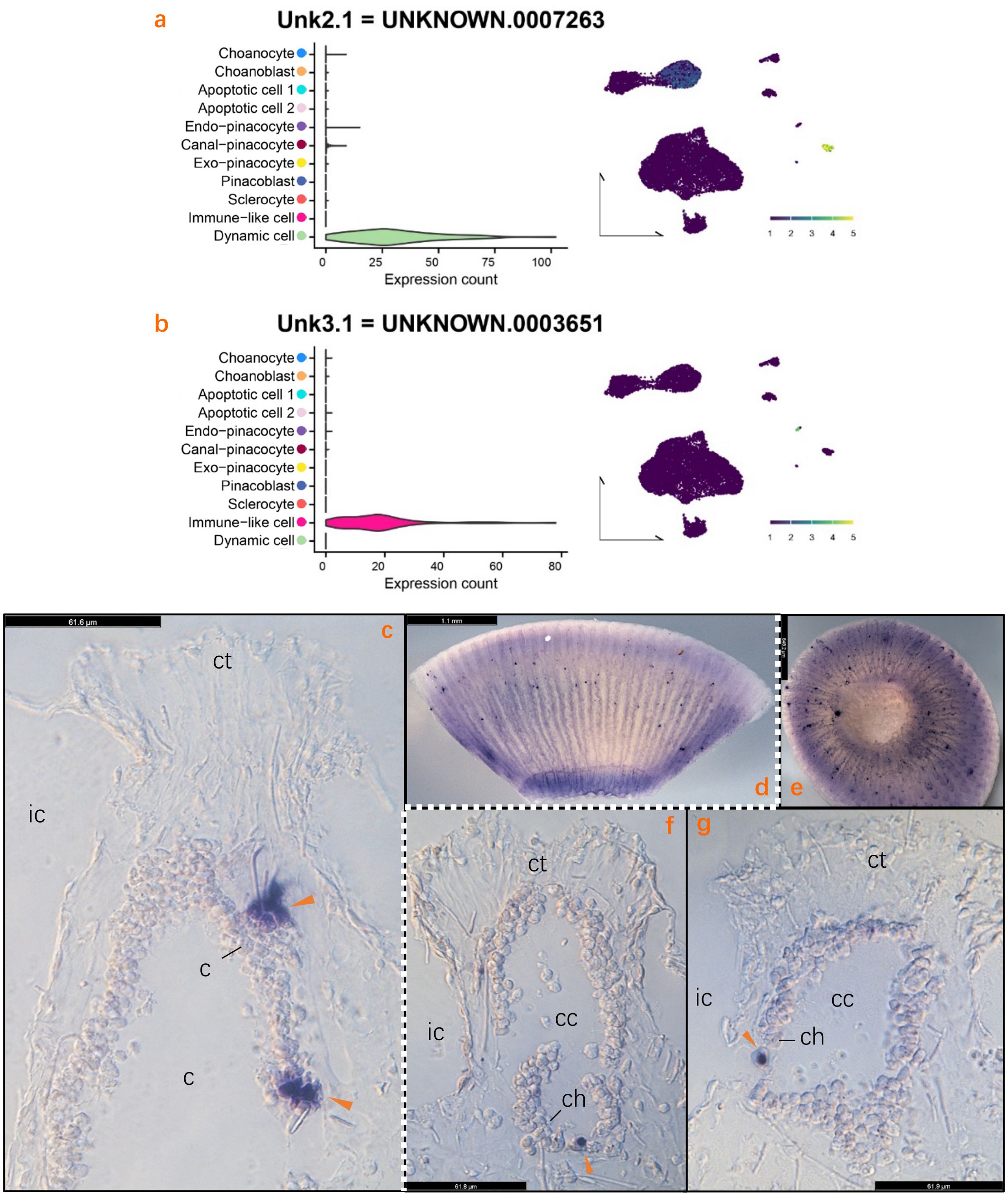


**figure S15. Gene expression visualization of the marker genes of dynamic cells and immune-like cells. a**, **c-d**, The expression level and cell distribution of the marker gene of dynamic cells; **b**, **e-g**, The expression level and cell distribution of the marker gene of immune-like cells. The purple colour in the hybridization results indicates that the cells at that location highly express the specified marker gene.

a (figure S16)

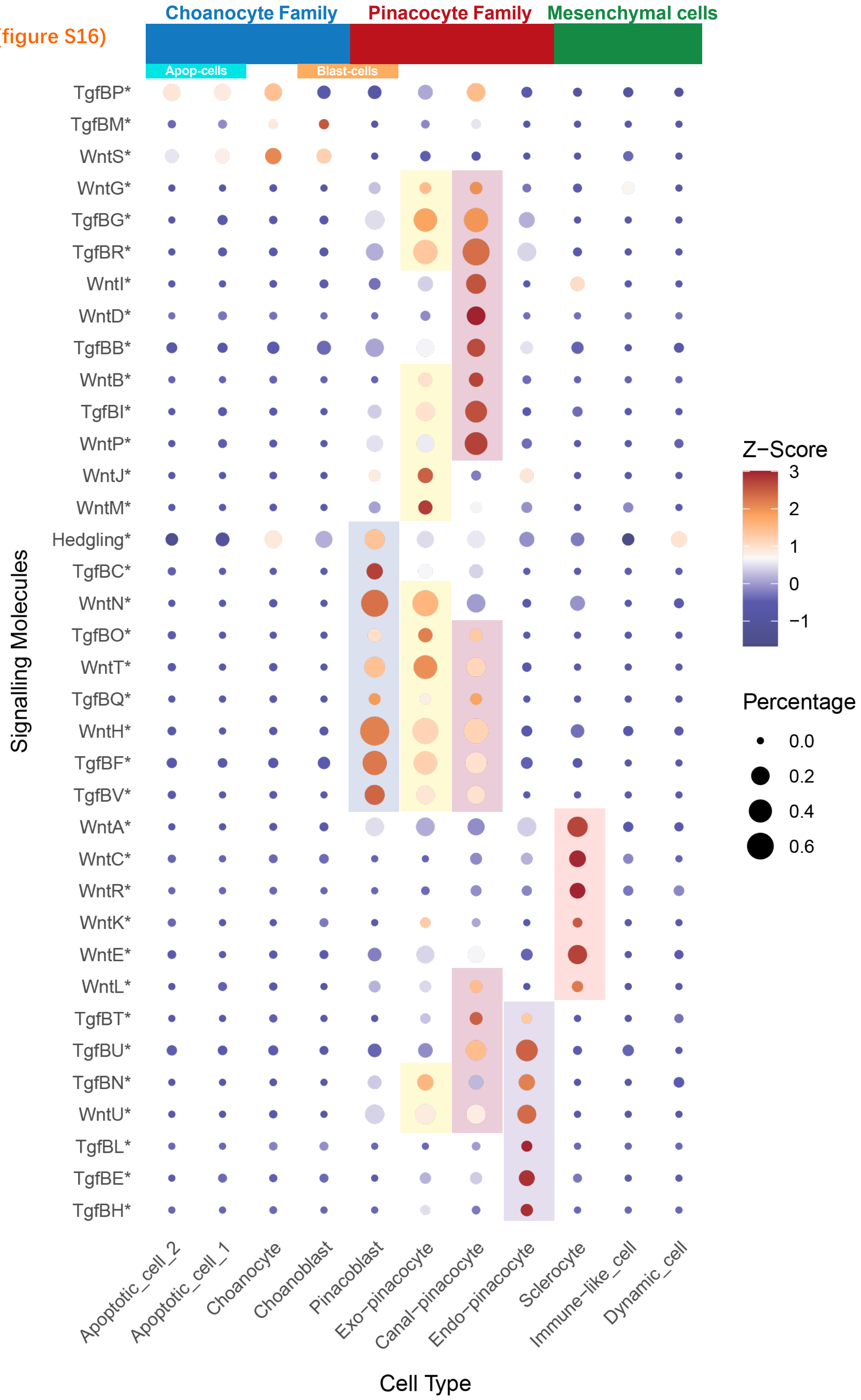
Choanocyte Family
Pinacocyte Family
Mesenchymal cells
Apop-cells
Blast-cells
TgfBP*
TgfBM*
WntS*
WntG*
TgfBG*
TgfBR*
WntI*
WntD*
TgfBB*
WntB*
TgfBI*
WntP*
WntJ*
WntM*
Hedgling*
TgfBC*
WntN*
TgfBO*
WntT*
TgfBQ*
WntH*
TgfBF*
TgfBV*
WntA*
WntC*
WntR*
WntK*
WntE*
WntL*
TgfBT*
TgfBU*
TgfBN*
WntU*
TgfBL*
TgfBE*
TgfBH*
Signalling Molecules
Apoptotic_cell_2
Apoptotic_cell_1
Choanocyte
Choanoblast
Pinacoblast
Exo−pinacocyte
Canal−pinacocyte
Endo−pinacocyte
Sclerocyte
Immune−like_cell
Dynamic_cell
Cell Type
Z−Score
3
2
1
0
−1
Percentage
0.0
0.2
0.4
0.6

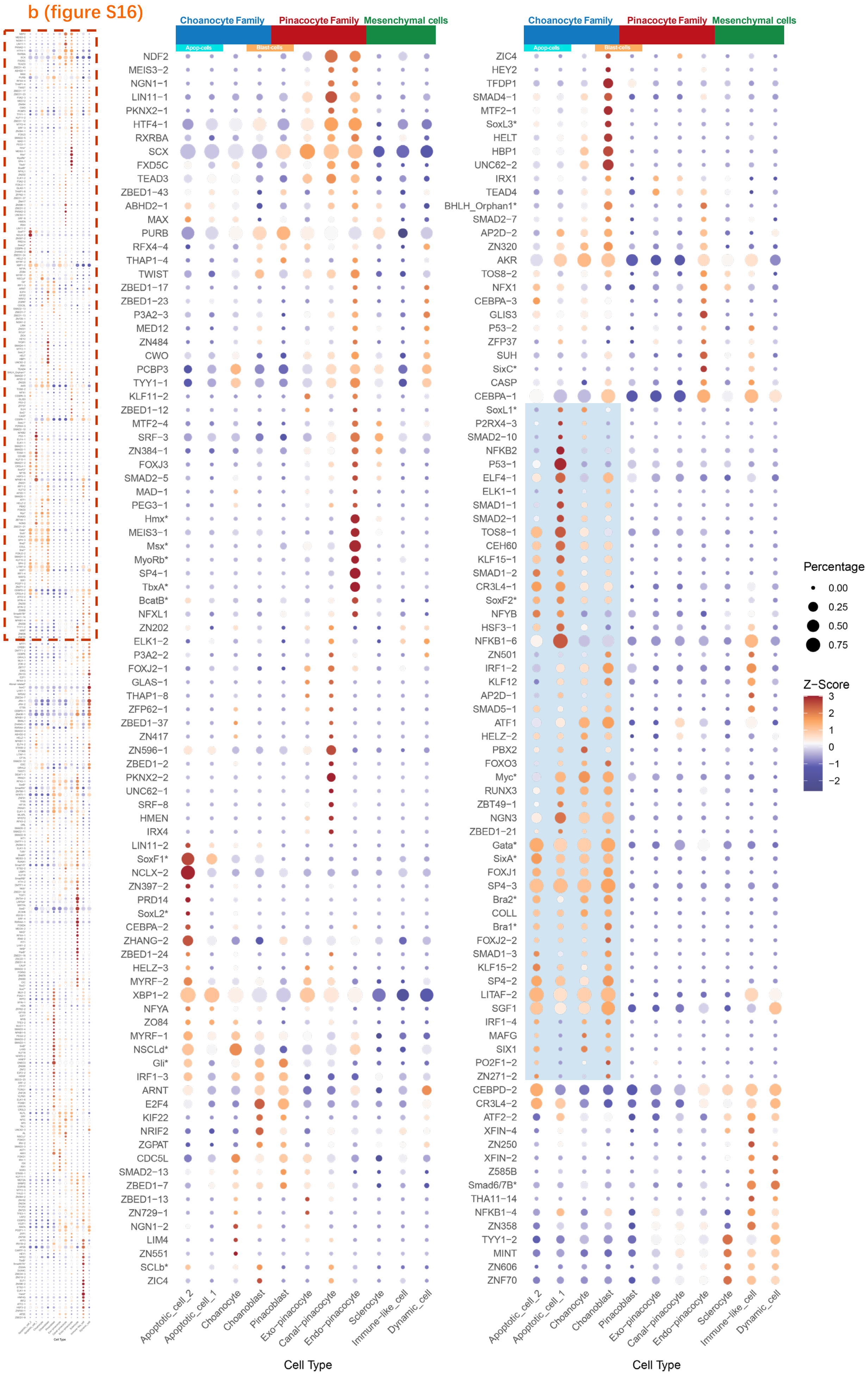

b (figure S16)
Choanocyte Family
Pinacocyte Family
Mesenchymal cells
Apop-cells
Blast-cells
NDF2
MEIS3−2
NGN1−1
LIN11−1
PKNX2−1
HTF4−1
RXRBA
SCX
FXD5C
TEAD3
ZBED1−43
ABHD2−1
MAX
PURB
RFX4−4
THAP1−4
TWIST
ZBED1−17
ZBED1−23
P3A2−3
MED12
ZN484
CWO
PCBP3
TYY1−1
KLF11−2
ZBED1−12
MTF2−4
SRF−3
ZN384−1
FOXJ3
SMAD2−5
MAD−1
PEG3−1
Hmx*
MEIS3−1
Msx*
MyoRb*
SP4−1
TbxA*
BcatB*
NFXL1
ZN202
ELK1−2
P3A2−2
FOXJ2−1
GLAS−1
THAP1−8
ZFP62−1
ZBED1−37
ZN417
ZN596−1
ZBED1−2
PKNX2−2
UNC62−1
SRF−8
HMEN
IRX4
LIN11−2
SoxF1*
NCLX−2
ZN397−2
PRD14
SoxL2*
CEBPA−2
ZHANG−2
ZBED1−24
HELZ−3
MYRF−2
XBP1−2
NFYA
ZO84
MYRF−1
NSCLd*
Gli*
IRF1−3
ARNT
E2F4
KIF22
NRIF2
ZGPAT
CDC5L
SMAD2−13
ZBED1−7
ZBED1−13
ZN729−1
NGN1−2
LIM4
ZN551
SCLb*
ZIC4
ZIC4
HEY2
TFDP1
SMAD4−1
MTF2−1
SoxL3*
HELT
HBP1
UNC62−2
IRX1
TEAD4
BHLH_Orphan1*
SMAD2−7
AP2D−2
ZN320
AKR
TOS8−2
NFX1
CEBPA−3
GLIS3
P53−2
ZFP37
SUH
SixC*
CASP
CEBPA−1
SoxL1*
P2RX4−3
SMAD2−10
NFKB2
P53−1
ELF4−1
ELK1−1
SMAD1−1
SMAD2−1
TOS8−1
CEH60
KLF15−1
SMAD1−2
CR3L4−1
SoxF2*
NFYB
HSF3−1
NFKB1−6
ZN501
IRF1−2
KLF12
AP2D−1
SMAD5−1
ATF1
HELZ−2
PBX2
FOXO3
Myc*
RUNX3
ZBT49−1
NGN3
ZBED1−21
Gata*
SixA*
FOXJ1
SP4−3
Bra2*
COLL
Bra1*
FOXJ2−2
SMAD1−3
KLF15−2
SP4−2
LITAF−2
SGF1
IRF1−4
MAFG
SIX1
PO2F1−2
ZN271−2
CEBPD−2
CR3L4−2
ATF2−2
XFIN−4
ZN250
XFIN−2
Z585B
Smad6/7B*
THA11−14
NFKB1−4
ZN358
TYY1−2
MINT
ZN606
ZNF70
Apoptotic_cell_2
Apoptotic_cell_1
Choanocyte
Choanoblast
Pinacoblast
Exo−pinacocyte
Canal−pinacocyte
Endo−pinacocyte
Sclerocyte
Immune−like_cell
Dynamic_cell
Cell Type
Percentage
0.00
0.25
0.50
0.75
Z−Score
3
2
1
0
−1
−2

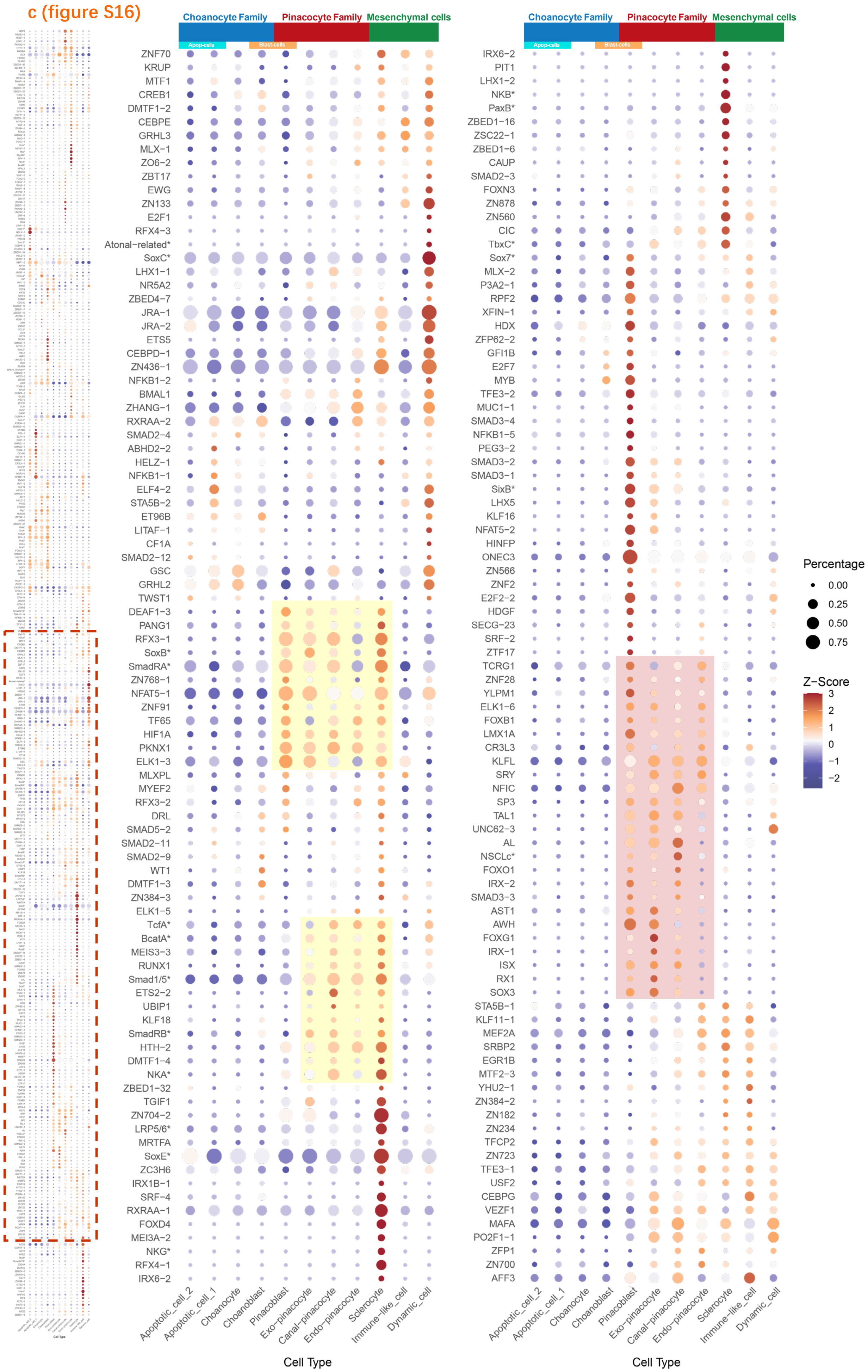
c (figure S16)
Choanocyte Family
Pinacocyte Family
Mesenchymal cells
Apop-cells
Blast-cells
Percentage
0.00
0.25
0.50
0.75
Z−Score
Cell Type
Apoptotic_cell_2
Apoptotic_cell_1
Choanocyte
Choanoblast
Pinacoblast
Exo−pinacocyte
Canal−pinacocyte
Endo−pinacocyte
Sclerocyte
Immune−like_cell
Dynamic_cell

d (figure S16)

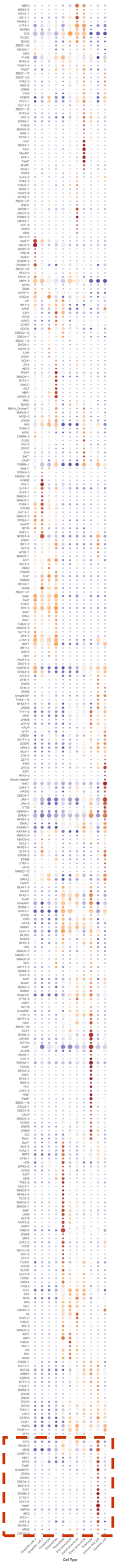

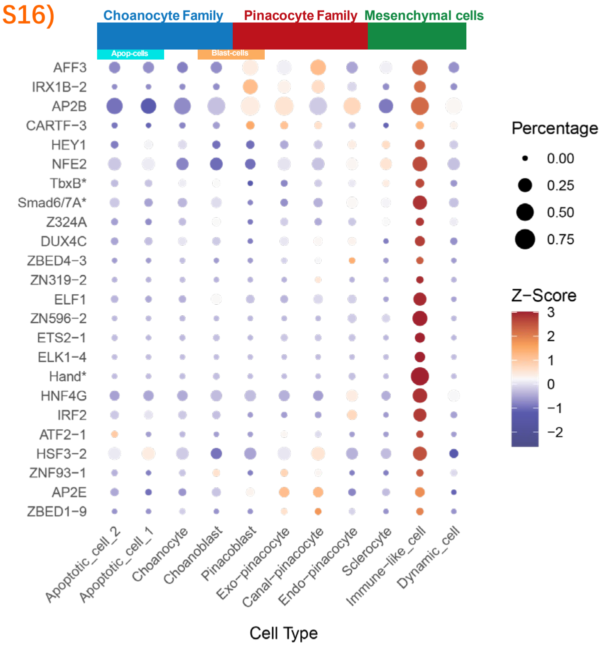


**figure S16. The expression of signalling molecules and transcription factors annotated by the GO term as well as the reciprocal best hit (RBH). a,** Dot plot showing expression of all *Wnt*, *Tgf,* and *Hedgling* that could be annotated in *S. capricorn*; **b-d,** Dot plot divided into three parts showing expression of all transcription factors that could be annotated in *S. capricorn*. Dot plot clusters according to the z-score values, and some clusters have been highlighted by colored rectangles. The dot's size indicates the gene's expression ratio in the corresponding type of cells, that is, the percentage. The dot's colour indicates the gene's expression intensity in different cell types, that is, the z-score. Red indicates high expression, and blue indicates low expression. The darker the colour, the more it deviates from the average expression. The annotations marked with an asterisk indicate transcription factors are the RBH results with the published sequence in *S. ciliatum*. Note that unlike all the other plots shown, the order of cell types in the choanocyte and pinacocyte families has been changed here, using blast cells as the connecting point.

**Appendix II (Reagents)**

- ACME with NaCl solution: 5M NaCl solution 801uL, Glycerol 5mL, Methanol (*Sigma*, 1.06009) 7.5mL, Acetic acid (*Sigma*, A6283) 5mL, add RNase-free water to 50mL (without light, RT storage);
- Ca/Mg-FASW 1L: NaCl 26.24g, KCl 0.671g, $Na_2SO_4$ 4.687g, $NaHCO_3$ 0.1806g, 1M Tris(pH8) 10mL, 0.5M EGTA(pH8) (*Bioworld*, 40520008) 5mL, add RO water to 1L;
- Hybridisation buffer (HB): Formamide 25mL, 20X Saline-sodium citrate 12.5mL, 50mg/mL Heparin (*Sigma*, H9399) 100uL, 0.5M EDTA(pH7) 500uL, 50x Denhardts filt (AMRESCO, E717-50ML) 1000uL, 10mg/mL yeast tRNA (*Sigma*, R8508) 500uL, 20% Tween-20 250uL, add RO water to 50mL;
- Mg-free AP buffer: 5M NaCl 1mL, 1M Tris(pH9.5) 5mL, 20% Tween-20 250uL, add RO water to 50mL;
- MOPS fix: 1M MOPS (pH7.5) (*Sigma*, M1254) 4.95mL, 5M NaCl 4.95mL, 1M $MgSO_4$ 0.1mL, 20% PFA (*Sigma*, 441244) 10mL, 25% Glutaraldehyde (*Sigma*, G5882) 100uL, 0.5M EDTA 0.5mL, add RO water to 50mL;
- 2% blocking buffer in maleic acid buffer (MAB): 1g Casein from bovine milk (*Sigma*, C5890), 1M Maleic acid(pH7.5) (*Sigma*, M0375) 5mL, 5M NaCl 1.25mL, 20% Tween-20 250uL, add RO water to 50mL;
- 10% sucrose in TBST (Tris-buffered saline with 0.1% Tween-20 detergent): 50g sucrose and 1 TBST tablet (*Sigma*, 91414) dissolved in 500mL of RO water.

### Appendix III (probe table)

| Probe name | Gene name | Cell type | Concent-ration for *in situ* | Annotation | Template sequence (including T3-pol sequence) |
|---|---|---|---|---|---|
| A.Cho3 | UNKNOWN.0025773 | Choanocyte family | 2.5ng/uL | scag4.0025773; UNKNOWN; unknow, signal peptide cleavage predicted [scahrs1_100 s.ca] | AATTAACCCTCACTAAAGCATACACTTTCTACTCAGACTTGGCAAACAAGACGTTCAGCTCGGTTCGTGTGAAAAGAAGGCTCTTATAATGAGGTGCTAATAATAATAACTTTAATGTGCAATTGGAACAAGATAGAAATCAATTGGGGAGGATACACTGTTCGGTGTAGGGAACACAAATAAGGTTAGCTATACAAACTATACAAGTATATCTGACAGAAAGAGACAAGCAGGGAGCAATGTTTCTCTTTGCTCTGAGTCCGTCTGCGGAGTGGTGTTGAAGATTAGTACATCTGAAAGATGGAAGAAGAGGCAGAGATCTAGAACGTATATTCGGATGTACAACGAGGAGCAACACAGCACAGTAGTTGCTCATGAAAACAAGTATTTACTCAATGCCGCTTGGGATCATATTTGATATCGACACGCCTCCGGTAGGTGTCACATCTTCCGCGGGTCTGATCCTCTCCAGCCGCGTTCCGGTAGGTGGAACAAAACCGGGCCATGATGGTCTTGGTACCGCAGTCATCATACTGATGCGTCTCCTGCACAGGCGTCAAGACGTCGGAGTAGCCGTGGGTCCTGGTGCCATCGTCCCAGTCAATGCACCACGAGTATCCCTGGTGGGTTTGAGTGCAGGTGCCGTACCAGGTGATCAGTGCAACGTTGGCACGACCTCCGGAGAAAGGCTCGATGACAGAGGCGTTCTTTTTCGTTGGGTTGTTCTTTGGATACTGGATGATTTGAGGCCTGCAGCCGTACTTGCAACCACATGTGAGTTCATCCATAGACACCATGTCCTCCTCCATTTGGCTGATGAAGTCGTTCACGTCCAGTTGAGCGCCGGCCATGGTGAACCCGGCCACGCAAAGGGCAAGAACGACGAGAACCTTCATTGCTGTTGCTGTCTTCCTATCGTGCGCGCACTGTACAAGGTACTATGGGAGGTCTACAGTAGGCAATGAAGAATCCCGAGCTGTGCTGCTGCTTTTATACCA |
| ChoX.2 | UNKNOWN.0003730 | Apoptotic cell 1 | 2.5ng/uL | scag4.0003730; UNKNOWN; unknow, predicted 1 transmembrane helices [scahrs1_3 s.ca] | AATTAACCCTCACTAAAGGGTGTGTTATGTGTGTTCCTACCTCAGAAATACATCATGCAAAGTTTGATTGAACTACAAGATAGCGCTTGAGCTGCAAGGCCTTGAAGAAAGTATTGAGATTGTTTTGGAATCTACTTTAGGACAGTTTAGGACAAGGCCCAATATACACCAGAGCTTTGCCAAGAAGAACCACTCTCCAGAAATCCAGGACTCCTGAACAAACCCACCAGGTAAGAATACTTCAGGAATATGACAAAAAAATGTAACGATAATAATTATGAAGGAATTCGTAAAGAATCACTGTAATATGCTTTTGCCTCCAGCTGTCATCTACTTAAATTAAGAGCATAACAACTTATTCCACAAGAACGAAAAGGACAAATATATTCTTTGGAAAGAACTGCTCAGCTCTGTGAGTAATCCACGGCACTACTGTATTATGGCAATTGAACAGGACCGTCAAGTCGAAATAGCAGAAACCTACTTCAAGCACTACATTGTCAAGTGTTTATTTACCATGCTAAACTACAATCGAATATACTCACGGGATTTGAAAAGGCAATAGTCTCACATAACATTAATACTATCCTAGACAACAAGCCCAGAAAATGAACTAAGAAGGGAGTTTCAACAAAAATCGAACCAATTGATGCATTCAGATCACTTCATATGGTAAACAACATTCAAGCTAGTATGTGTACTACTGTAAAGGTTTATTTTATCAAGTACCATTCAGCGATCTCCAATCGATCTCTCACCAGGTGTCTTCAGAAATGTGTTGGCTGACTACGACGTTGATGGAAGTGTCGGCGCCATTCTCACGAGATGCGTCTTCTTTAGCCTGACGGCTGAAAAAAGCGGCATCCCGTAGGAACATCTCGCTTTGCACGGACCGCGGACAACTGGAGCGCAGGAGG |
| Cho3.3 | DNLI3-LIKE.0052832 | Apoptotic cell 2 | 3.27ng/uL | scag4.0052832; DNLI3_LIKE; DNA ligase 3 like (0 Homo) [scahrs1_348 s.ca] | AATTAACCCTCACTAAAGTCTGGCAGACCAATATCAGTTGCATTAGCAGCATACTCCTCGGGTAAGGTGAGTAATGAGTACTCAGTTCAGGGCTGTCACTGTTCTGTGCTTACAACGCTGATGAAGCCATGCAGCAAGCCAGCAAGCCAGCATGCACTTTGGTAATAACTTATACATGTCATTATTGTGTGGAGCAACTCGGACAAGCCAGACCACCTGACATTAGACTTTCTATTAGTTCTAATGCAAGTAGGCTCGATGACACAATACAGAAACGGGCACAATGCTGTGCACTAAAATTCACTTGAAATTAGCAAAAGCACGAAACGTTACAAAGTGCTGCGGCTCAGGCTAGGGTACAACCAAGCAAGTGGTTGTCAAGATGTGTTCTGCTCTGGAAGAAACCAGATTTACAGTGCAGCATCCAGTGATTACAAAGTGACAAAGCAATAATAAAAAGAAACAAAAATTAAAAACAAAAGACAATTGTGGCGTTCAATAAATTATCATGATCGTACCTTTGAAGTAAGTATTATTTCCTGCTAAATTCGATAAAAACATAATTAGGGCTCAAACGAAGCTTCGTGCTTAGCAAACAAGCCATGGATGAAGACATGCACCACAGCAATGCATGCTAAAAATTGAAGGTATTATGAAAGTTGAAGGTGAATGCACCATTGCGTAACGCATCAAGACGTGGCAGCATTGGACTGCACTGAAACTAGGATCTTTTTTTTTATAAAGGTTACATGGTTTCCTTGCATGACAGAAAATATAACAGCTCTAGCTAATATTTTTTGGAAATTCTGATAGAAAAGGATTCATTCTGGATTTCATGACATTAAAAAAAAAAAGCGTTTCAAACATCTACCTACCTACACTAGTTTTCTTGAACAAGATACACAATTATCGTTGCCATAGCAGAAGTGATGAATATTTCAAAGCTGTGATACTGGTTAGGAGAGGAGCAGGGGTAGGAGGTATGTCATATTATTGAC |
| Cho4.1 | H1E-LIKE.0025440 | Blast cell | 2.5ng/uL | scag4.0025440; H1E_like; Histone H1E like (9.69e-10 Chironomus) [scahrs1_100 s.ca] | AATTAACCCTCACTAAAGTGGGCCCTAAGAATAAACTCATGTATATAAGACGTACATCCAATGAGGGAAGGGGAAGTTGGGGAATAAAGAAAACGGCCGCTTTGATATTGAAATGCAGGGACAGCCATGTCTTTTCATGTGTCTGTGGTACACAAACTTACAAAGCTTTTTGCTTTAAAAAGCCCCTCGGTCAGTGCAATCTGATGCGGTGAATAATCATTTCTGTTCTTCACAAGGCCACCCCCAAACTGGGCCAAAGTATGTTCCCTTTCCCTTCATCAAAGGGAACAAGGCCAGAAACAGCAATCAACTGATTTTGCAGGAGAATTTGTGTTTCTTCAAAAATCAATGGCAAAAGATCTAGTAGCCAGAGAAGAGATTGACACAAATTAAAGAAAGCATGAGGCATAATCCGGTAGATTTATTCCATTGAAAACAAAAACACATCGAAACAAGAACAACAAGACTCAATAGCAGTCAAGTAGTTATATTGATAAAGGTACATCATATTGAGCTAAGATTCAGAAAAATCAAGACGCATTGCATTGAAAAACGCAGGGTAATAGTTCAGCAAATAAAGAAAGAAGCCAACATAACAGCACAGAATAAGTACAATATGGCGCAGAGAATTTTCAAGCTATGCACTATTGTTGTGAGTTGACTATGATTTGCCGGAGGAAAGATATGAAGACATGCGAAGTTTTGCCCGGTTTTTACATGAATATTTTGACCTATGGATGAGAGAATGTGAAGGTTTTATGCACTAAAGATGACTAAAAGAAACGAGCACGCTTACTTCTTCGGGGACTTGGGTACCTTCTTCGCTGCTGGCTTCTTGGCGGCTGGT |
| Unk1.1 | YJ48-LIKE.0054519 | Endo-pinacocyte | 1.5ng/uL | scag4.0054519; YJ48_LIKE; Uncharacterized methyltransferase C70.08c like (4.93e-10 Schizosaccharomyces) [scahrs1_348 s.ca] | AATTAACCCTCACTAAAGATCTGTAGGGAAGGTCACACGAAATACGTGACACTGATGATAATGAAGTGCGAAAATTTACAATAGGGAAGTTACGGAAAGGGACAAATGGTGGAGCTATAGCGCTGGTTTGACGGCGACGATCGTAAGCATGGAGCAATACATACGCATCTTGGCATCCAAGCTTCCCAACTCGTGCTTAGCCAGGAAATCTTGCAACTCCCTCTCGGTCAATTTCCCAGTGTCAATAGCACCTTCAGTGTATGCCTCTAAGAAGTCGAAATATTCCCTGACGCTTGCAAACTTCAGTTGCATTTCGTCGTTCTGGACGGCGGAATGCACGTGAAGGCCTGCTTCGCTGCACAGGCCTCTCCACGAATCCAGCGATCGGTAGCGCATTCCGGTGACCTCCTCTTCACTCTTCCCCGTGATCAGAAGAGAGAGATCGTTGGCATATGGACCAGTGCCGAACGTAATGTTTGCGCAGAGACGACCTTCAGGCTTCAGACACCTCGCCACACTGATCATGGTGGCGCTGATATCATCTTCCTCGACCCAGTGCAGTGCGAAGTTCGAGAACACCGCGTCAAACGGCCCATGATCCACAGCATCGTAAACAACACCTGTGATAAACGTCACATTACTTCCACCATCGTTAGGAGACGATTCGTTCTTGGTCTCCATCCTTGCCGCATCCAGCTGGGCTCTTTTGATACGCTTGACGTTTGGATCAACGCCCAGGACACTTCCCTGACGTCCCACGGCTGATCCTATTTCCATTGCCAATTCACCGGTCCCGCATCCCAAGTCCAGGATCCTGTCACCGGTCTGTGGACGGAGTTTCGATCCAACGAGAAGCTGTCCGTACTGGTGCTGTACACTGGAATGGTCTTCATAGAACTGAGCAAGGAAGGACGATGCCTTCCGCTGTAACTTCGGCATCTTTCCCCTCCCAGACATATTGCGGGCAGACACGAATGTATTATGAA |
| Pin1.9 | USH2A-LIKE.0031340 | Canal-pinacocyte | 2.5ng/uL | scag4.0031340; USH2A_LIKE; Usherin like (1.08e-102 Mus); signal peptide cleavage predicted [scahrs1_100 s.ca] | AATTAACCCTCACTAAAGGATGCAAATGCCACTCCCTTCTGAGCAGTTTCCGGATGCAAACTGCTGGCCGGCAAGATTGCAGCCACATTCTGTGCAATTGACAGTGAGGTCAATATTCATCGGATCTCCAGGATGGAAGAACTGAGGAACACATTGCTGGCAGTTTATTCCGGTTGTGAGACCCTGGCATGCAACGCACACGCCACCACCGGACAAGACATTGCCAGTATTGAGGCTCCTTCCAACGGCACTGGGGTCGTACACGCAGGATTCAGCATGTCCGAAGCACTGGCATTGCTCGCAGGCGACTCCAGACATGAATGGCTGTTGGTTGAAAAGTGGCTGACACATCTCACAGTTGACACCCTGGACGAAGCGAGTAGGCTCGCACATGCACACGTAACCGCCGTTTTGCGCCAGCTCACAGTCAGTAGCATGTCCATTGCATATGCAATGACCTGCCACGCTCATAAGCTGAATTGTGTAGAATGGAAAAGCAGGATCAGAGAAGCCTCTGAAACTAAGACGAACTGCATTAGCCGTGAAGAGGTCCACTACTGCTGGGTTGTTGCGAGGATTTATACGCATGCCATTTACAATGGCTGTTCTCTCGTTAAAGAGACGGAGTTCAAAGAATCGATCTGAAATGGAATTGATGTCCATATAGGTGATGCAGAATGGCGTGCCAATATTGGACCCGTCCAATAGCACAAAAGCCTGGGATGGAACCCCGAAGATGGATTGGCACTGAGACTCCGTTGATACCACGTAGATGAATGGAGTGAACCCATTGCCGTGGTCACCACTGAGGAGCAGGGCCTCCGGAGGCGGCGACGACATGGTCACTTGGATGAAATTGACAACGAATGGCTGTTCGAATTGCACTGTCAATGAGGCATTGACAAGATTATTGGCTGACTTCCAAAAGGTATCTTCTGCCGCTGTACCTGCACTGGGAGGCAGGTTACTGGTCAGGTTGCACA |
| Pin2.1 | UNKNOWN.0010448 | Exo-pinacocyte | 0.5ng/uL | scag4.0010448; UNKNOWN; unknow, signal peptide cleavage predicted [scahrs1_40 s.ca] | AATTAACCCTCACTAAAGGAGACTTCTGCAATAAAGCAATGCACTGCCTACTTTCTAGCATGCTGCGGGAGGCACAAGGAGGACTGCATGCCACACCGCTGCAGAATGCCAAATTAGGTACCATAGCAAGGCCTGGACGGACACAGACGCCTCACCTCCTAACCGCGATAATAGGGCCTGTAATAGGGCCTGGGTGGTGGAGGCCTGTAGTAGCGAATACGTGGCAGGCGGAACCTTGGCAGACGTATGCGCGGCAGACGGATGCGAGGTAGACGGATGCGAGGTAGACGAATGCGGGGCAAGCGAATTCGAGGCAGGCGGATACGTGGGATGCGGATGCGTTTGACACGTCTAATGAGTCGTCGAAGGAATGGAGAGCGACCGTTGACGGCCTCATTTTCTTGTAGTGCAGCATACTCGACATCATCCTGTCCGACGTCGTCCTCCTCGTCATCCTCGTCATCGTCCATGACCAACTGGACATCGCCATCCCGAGCGTAATCACTGTCGTACTCAAGATCAGCATTACTGTCGTCTC |
| Scl.1 | UNKNOWN.0022356 | Sclerocyte | 0.6ng/uL | scag4.0022356; UNKNOWN; unknow, predicted 1 transmembrane helices, signal peptide cleavage predicted [scahrs1_100 s.ca] | AATTAACCCTCACTAAAGTCAGTGAGAGGGAGATCAGTGAGGGATCAGGCAGAATTGAGCAAGTGATCAGCGAGACTGGATTGGAGGGATAGTTGAGTCGAACAGTAGTGAGGTGAAGTGGAACTACTTACGGGCAAGCCTCGGATCAAAGGGAAATTGGAAGCCTAGAAAACAACTAGTGTGGTGGGCTGAACATCAGTGCTGACAGAGTTACGGCATGAAGATCGCCACAAGGGCCATGGCGAGGACGGAGAAGAGGCTGATGGTCAGCCGAACGCCGCTGGCGCGCGCATCAATGACGTTCTGCAGCTCATCGCAACAGTTGTCGCTGATGGTGGCGTTATAGTCGCACTGCTGCCATTGCTGCGCGGCGTAGGGCAGGATGACACAGCAGGCCTCGTCGTCGGCTTTGTGAGCCAGGTGCTCTGCGTTCAGCGTCACCGTACCAGTGTTGACAGCGTTCATAGGGGTGGTCTGTCTCTTCGTGCGATCCTTCACGAAGTTCTTGATCTTGCTGGTGAAGCCCTTAAGGAAACCACCAATGGACGAGTCAGCCTTGCACTCACCCAGCTCGGTGGACTCCGATGTCCACATCACAGTAAATGTGATGTTTCCCCCCGCCGCCATCCACGAAATCGTACTCGGAGTCATAGAAGAGTTATCGGCGGTCATGCACTTTGACTCGTCGACGTTGCAAGGCATTTTCACTTCGTGATGCTCCTCGTGGTCGTGATCGTCACCGTCTTCCCCGTGATCGTGGCCACCGCTTTCTTCCCTGACGTGCCAATGATCTTGGGACGCCTGTGCCAGCTGCGTCAGCTGCAGCAAGAGCACGGCCAGTGCGGATATCATCAGGACCTTCATTATGACACTAGAGAGCCGAGTTGCTACCTGTCCCGCACAAGATTACCGACAA |
| Unk2.1 | UNKNOWN.0007263 | Dynamic cell | 2.5ng/uL | scag4.0007263; UNKNOWN; unknow [scahrs1_11 s.ca] | AATTAACCCTCACTAAAGGCTCAGTAACTAGTTACTGCTAAACAAGTTTTTCCAACCGTCTATTTTGTGCCTACCTGTGTGTGTTCACGCAAGTGTTGGCTTGTAGGCAAGTTATCTCGAACCCCTGGCGGAAGTATTCGTTCATCTTTTAATTTATCACAGTCCAGAGGCTGTCGGTTGCATGTTCGTGGGGACGTTTTATGAACACAAGGGTTGGTAATAATGCTTGAGATACAGAATAGCAAAGATACTCCACGTGTGTTTACCTTATAACGTGAACAAATTGGCCCCCAGTCTTGGGCAAGGTCTTGGGCAAGGTACATGTGCCTGTCTACTTGTAGTGCAGAGATTTCCCCTACAATGTTGTTACATCATACAGTGCTTCGGGGTATTTTTCATGATATCTGGTGCACACGTGATGCGAGTGTATGACCTTGGTTCATCAGCCTGTCTTCTCCAAAAAGAAGTAGCTAGTTGTATGCTTCCAAGTGGGGATGGGGTGTTTTCACGCGCACGATGG |
